# A novel zebrafish intestinal tumor model reveals a role for *cyp7a1*-dependent tumor-liver crosstalk in tumor's adverse effects on host

**DOI:** 10.1101/199349

**Authors:** Sora Enya, Koichi Kawakami, Yutaka Suzuki, Shinpei Kawaoka

## Abstract

The nature of host organs and genes that underlie tumor-induced physiological disruption on host remains ill-defined. Here, we establish a novel zebrafish intestinal tumor model that is optimized for addressing this issue, and find that hepatic *cyp7a1*, the rate-limiting factor for synthesizing bile acids (BAs), is such a host gene. Inducing *kras*^*G12D*^ by *Gal4* specifically expressed in the posterior intestine resulted in formation of an intestinal tumor classified as dysplasia. The local intestinal tumor caused systemic detrimental effects on host including liver inflammation, hepatomegaly, growth defects, and organismal death. Whole-organismal level gene expression analysis and metabolite measurements revealed that the intestinal tumor reduced total BAs levels via down-regulation of hepatic *cyp7a1*. Genetically rescuing *cyp7a1* expression in the liver restored the BAs synthesis and ameliorated tumor-induced liver inflammation, but not other tumor-dependent phenotypes. Thus, we found a previously unknown role of *cyp7a1* as the host gene that links the intestinal tumor, hepatic cholesterol-BAs metabolism, and liver inflammation in tumor-bearing fish. Our model provides an important basis to discover host genes responsible for tumor-induced phenotypes and to uncover mechanisms underlying how tumors adversely affect host organisms.

## Introduction

Tumors disrupt host physiology in various ways, ultimately leading to organismal death (Egeblad et al., 2010; Fearon et al., 2012; McAllister and Weinberg, 2014; Owusu-Ansah and Perrimon, 2015). Mechanisms underlying physiological disruption by tumors involve inter-organ communication between tumors and normal organs. Due to its complex nature, how tumors affect host organs—and when and how host organs detect and respond to tumors— have remained largely elusive. In particular, host genes and signaling cascades mediating tumor-organ interaction (and thus tumor-induced phenotypes) are poorly defined. Understanding the nature of tumor-organ interaction and its mediator(s) at the genetic level is essential to understand how tumors interfere with host physiology, and to suggest a therapy that buffers tumor-dependent physiological disruption on host.

Animal models that are amenable to whole-organismal level experiments and genetic manipulations provide a tool for discovering physiologically important tumor-organ interaction and underlying mechanisms behind them. The fly *Drosophila melanogaster* is one such model. A fly tumor originating from the eye imaginal disc secretes insulin-like peptide 8 (Dilp8) to delay organismal growth and maturation, thereby enabling, or forcing, the organism to coordinate their overall growth with a local disease state (Garelli et al., 2012). Consistent with local disrupted states having influence on distant processes such as growth, physiological disruption such as wounding also induces a Dilp8-dependent growth delay (Colombani et al., 2015; Colombani et al., 2012; Garelli et al., 2012; Garelli et al., 2015; Katsuyama et al., 2015; Owusu-Ansah and Perrimon, 2015; Vallejo et al., 2015). In this phenomenon, Lgr3, the receptor for Dilp8 expressed in neurons, is the host protein responsible for the tumor-dependent growth delay (Colombani et al., 2015; Garelli et al., 2015; Vallejo et al., 2015). These studies establish the concept that organisms are able to sense local physiological disruption that can be spread systemically (Owusu-Ansah and Perrimon, 2015). Others have shown that fly tumors produce ImpL2, an antagonist for insulin-like growth factors, to cause loss of peripheral tissues including muscle and fat, a phenomenon called cachexia (Fearon et al., 2012; Figueroa-Clarevega and Bilder, 2015;Kwon et al., 2015). Such hormone-mediated mechanisms of cancer-induced cachexia have also been reported also in mice. For example, lung cancer secretes parathyroid-related hormone (PTHrP) that increases fat thermogenesis through its receptor PTHR, a host gene expressed in fat cells, resulting in cachexia (Kir et al., 2016; Kir et al., 2014). In another example, adipose triglyceride lipases have been implicated in cachexia, since mice lacking these lipases become resistant to cancer-induced fat loss (Das et al., 2011). In addition, tumors often elicit massive inflammation in distant organs, which is thought to affect whole-organismal physiology (Egeblad et al., 2010; Fearon et al., 2012; McAllister and Weinberg, 2014; Owusu-Ansah and Perrimon, 2015). These tumor-induced phenomena are highly heterogeneous: the same tumors do not always cause the same systemic phenotypes (Fearon et al., 2012). This indicates that these phenotypes are influenced by host genotype and physiology, and *vice versa*, and thus appear to behave in a context-dependent manner. Most importantly, as described above, even in cachexia, a well-known tumor-induced phenotype, only a small set of host genes responsible for this phenomenon have been identified.

Zebrafish is an emerging model for studying tumors (White et al., 2013) and tumor-organ interaction due to its plethora of advantages including (i) they are a vertebrate that gives rise to numerous offspring at once, (ii) larvae are transparent, enabling researchers to observe tumorigenesis and tumor-induced phenotypes easily in live animals, (iii) they are small enough to allow whole-organismal level experiments, and (iv) genetic manipulations are relatively easy and affordable when compared especially to mice. As a good example, zebrafish melanoma models have provided various insights into melanoma development in vivo (Kaufman et al., 2016; Lister et al., 2014; Santoriello et al., 2010; White et al., 2011). Zebrafish genetic tumor models currently available often develop tumors at relatively later stages of zebrafish development, mostly after pigmentation (White et al., 2013). In such cases, it takes time (several weeks to months) to obtain tumor-bearing fish, and they are already opaque when tumors arise unless the *casper* mutation is introduced (White et al., 2008). Hence, it would be meaningful to create a novel zebrafish tumor model where tumor formation and proliferation occur in the transparent stage of zebrafish development.

Furthermore, as is the case for zebrafish, most animal tumor models develop tumors at an adult stage, thereby preventing us from investigation into how tumors affect growing, juvenile vertebrates. For these reasons, a novel zebrafish tumor model is required.

In the current study, we successfully generated a novel intestinal tumor model. Careful characterization of this model led to the identification of four tumor-induced phenotypes including systemic inflammation, hepatomegaly, growth defects, and organismal death, which are seen even in human cancer patients. Anomalies in gene expression and metabolism were found in both the intestinal tumor and the distant liver upon whole-organismal level transcriptome analysis. On the basis of these, we found that a tumor-liver crosstalk, which can be defined by reduced expression of hepatic *cyp7a1* accompanied with altered cholesterol-bile acids flux, promote infiltration of neutrophils to the liver (liver inflammation) in tumor-bearing fish.

## Results

### pInt-Gal4-driven kras^*G12D*^ expression causes outgrowth of posterior intestine, leading to formation of the intestinal tumor

In order to generate a zebrafish model of tumorigenesis with early onset, we sought for *Gal4* line(s) capable of driving gene expression to a single organ (ie. organ specificity) at an early stage of zebrafish development. To this end, we crossed a set of *Gal4* lines (Asakawa and Kawakami, 2008; Asakawa et al., 2008) with a line *Tg(5×UAS:EGFP-P2A-kras*^*G12D*^) generated in this study with the Tol2 system (Fig. 1A and Table S1) (Kawakami, 2004; Kawakami et al., 1998)*. Tg(5×UAS:EGFP-P2A-kras*^*G12D*^) harbored a mutated *kras*, *kras*^*G12D*^, one of the most prevalent driver oncogenes in human malignant tumors (Fig. 1A and Table S1) (Schubbert et al., 2007). Expression of *kras*^*G12D*^ was linked with *EGFP* by *P2A*, a self-cleaving peptide sequence (Kim et al., 2011). Tissue outgrowth of *kras*^*G12D*^-positive cells was examined using fluorescence stereoscopic microscope within approximately 48 hours after observation of *Gal4*-dependent EGFP expression in a target organ.

**Figure 1.**
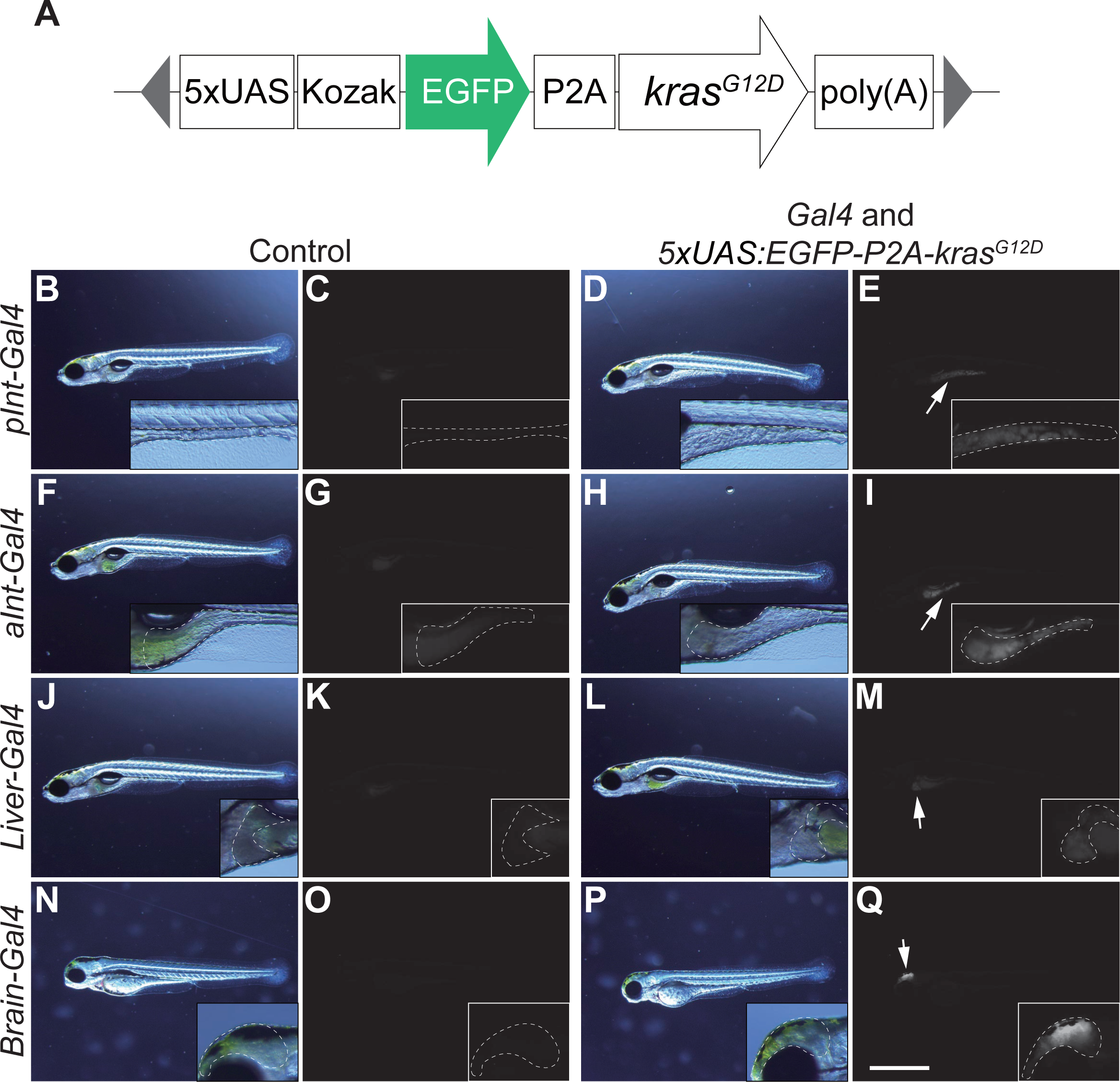
Screening a combination of *Gal4* lines and *5×UAS:EGFP-kras*^*G12D*^ transgene that causes outgrowth of target organs. (A) The structure of *5×UAS:EGFP-P2A-kras*^*G12D*^. The gray triangles represent the sequence recognized by Tol2 transposases. (B)-(Q) Screening for a *Gal4* line that is potent to induce outgrowth of target organs. Images of the sibling control (left) and *EGFP-kras*^*G12D*^-expressing fish (right) are shown. Higher-magnification images are also presented. (B), (D), (F), (H), (J), (L), (N), and (P) are bright filed images while the others are fluorescence images (EGFP). Target organs are outlined by white dots ((B)-(E) for gSAIzGFFD1105A (*pInt-Gal4*) (7 dpf), (F)-(I) for gSAIzGFFM103B (*aInt-Gal4*) (7 dpf), (J)-(M) for gSAIzGFFD886A (*Liver-Gal4*) (7 dpf), and (N)-(Q) for gSAGFF138A (*Brain-Gal4*) (3 dpf)). Fish without EGFP expression from the same clutch were used as sibling controls. White arrows indicate organs that express the *EGFP-kras*^*G12D*^ transgene. Scale bar indicates 1 mm.

Lines were identified showing the requisite expression in posterior intestinal cells (*pInt-Gal4*), anterior intestinal cells (*aInt-Gal4*), brain (*Brain-Gal4*), and liver (*Liver-Gal4*) (Fig. 1). From these, *pInt-Gal4* was chosen for further characterization due to its ability to cause efficient outgrowth of posterior intestinal cells upon *kras*^*G12D*^ expression (Fig. 1B-1E). *aInt-Gal4* was also able to cause outgrowth of anterior intestinal cells (Figs. 1F-1I). However, outgrowth of intestinal cells by *aInt-Gal4* was less dramatic when compared to that by *pInt-Gal4*. Moreover, expression of *aInt-Gal4*, despite specific after 5 dpf, was somewhat non-specific during 2-4 dpf, leading to abnormal growth of epidermal cells in a temporal manner (Fig. S1A-1D).

*pInt-Gal4* expression judged by EGFP expression was detectable from 4 dpf (days post-fertilization) ∼ 5 dpf (Fig. 2A-2B). Outgrowth of posterior intestinal cells by *pInt-Gal4*-driven *kras*^*G12D*^ expression was evident at 5 dpf (Fig. 2A-2B). Oncogene expression was confirmed by qPCR (Fig. 2C and Table S1). Moreover, 100% of fish harboring both *pInt-Gal4* and *5×UAS:EGFP-P2A-kras*^*G12D*^ exhibited the outgrowth phenotype at 5 dpf (Fig. S2A). Thus, at this stage, we were able to phenotypically discriminate tumor-bearing fish. The number of intestinal cells determined by DAPI-staining in *kras*^*G12D*^-expressing fish was significantly increased compared to that in the controls expressing EGFP under the regulation by *pInt-Gal4* (Fig. 2D-2J). In the previous study, Wallace et al. show that the mitotic rate of intestinal epithelial cells is high (∼ 40%) through 3 dpf, dropping at 4 ∼ 5 dpf (< 5 %) (Wallace et al., 2005). Despite the assumption that the majority of intestinal cells are post-mitotic at 5 dpf, we counted the number of mitotic cells by pH3 (phosphorylated histone H3)-staining (Fig. 2K-2S) and BrdU-incorporation experiments at this time point (Fig. S2B-S2J). The number of pH3-positive mitotic cells (Fig. 2K-2S) and BrdU-incorporated cells (Fig. S2B-S2J) were consistently higher in *kras*^*G12D*^-expressing fish than in the sibling controls, strongly suggesting that *pInt-Gal4*-driven *kras*^*G12D*^ expression promoted mitosis of intestinal cells.

**Figure 2.**
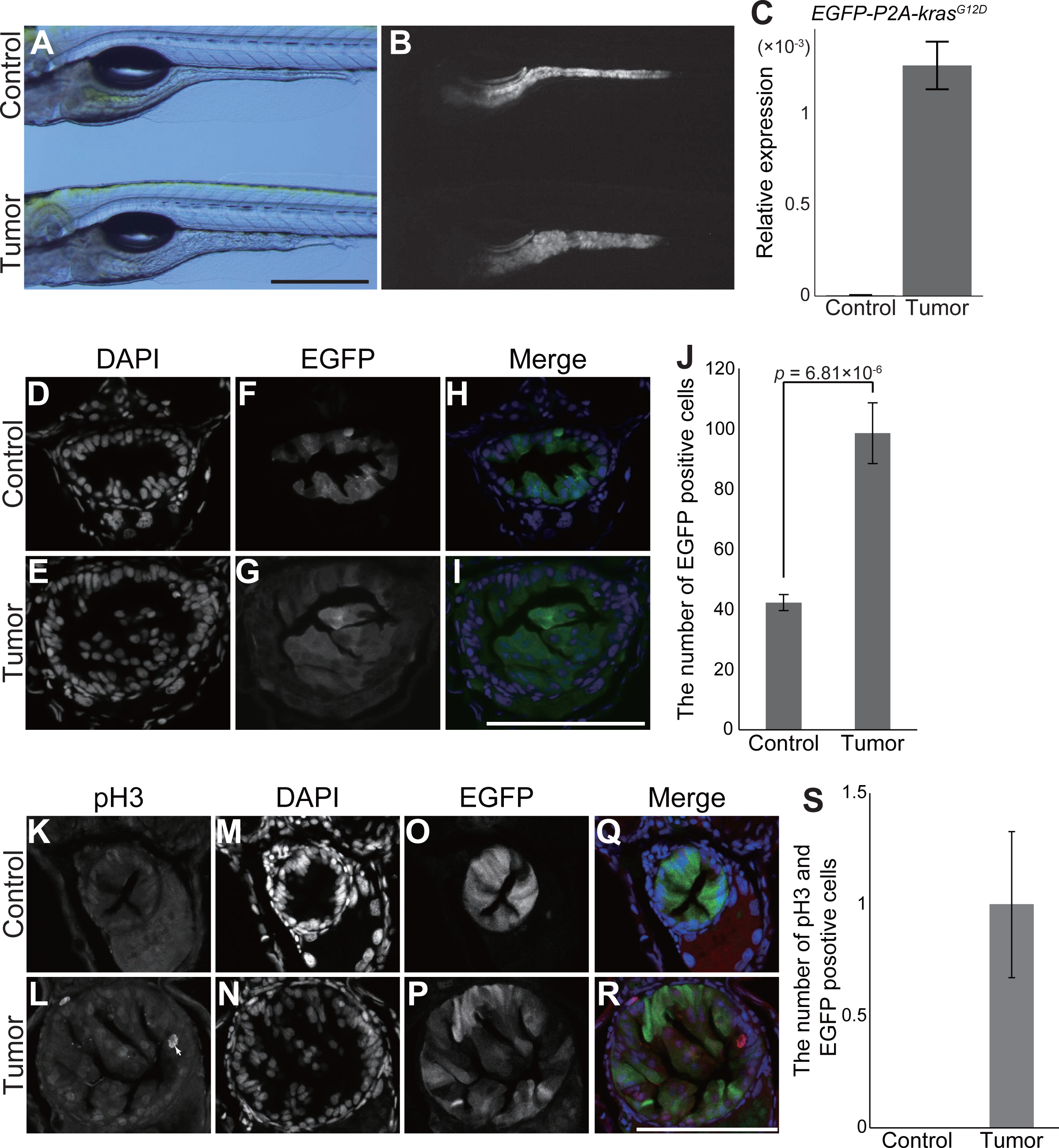
*pInt-Gal4* driven *kras*^*G12D*^ expression leads to abnormal proliferation of intestinal cells. (A)-(B) Representative images of tumor-bearing fish (*Tg(pInt-Gal4)*^*+/Tg*^*; Tg(5×UAS:EGFP-P2A-kras*^*G12D*^)^*+/Tg*^) and the sibling control (*Tg(pInt-Gal4)*^*+/Tg*^*; Tg(UAS:EGFP)*^*+/Tg*^) at 5 dpf. Bright field (A) and EGFP (B) images are shown. Scale bar indicates 500 μm. (C) qPCR for *EGFP-P2A-kras*^*G12D*^ expression in the sibling controls and tumor-bearing fish.The scores are normalized to expression of *rpl13a*. The data harbors three biological replicates. Error bars represent ± s.e.m. (D)-(I) Representative images of DAPI staining in intestine sections of tumor-bearing fish and the sibling controls at 5 dpf. DAPI (D, E) and EGFP (F, G) images are shown. In the merged images (H, I), DAPI and EGFP signals are shown in blue and green, respectively. Scale bar indicates 100 μm. (J) The number of EGFP and DAPI positive intestinal cells. The number of nuclei was manually counted from single section per individual fish. The data harbors 7 and 11 biological replicates from tumor-bearing fish and the sibling controls, respectively. Error bars represent ± s.e.m. Statistical significance was tested using student’s *t*-test (unpaired, one-tailed).(K)-(R) Representative images of fluorescent immunohistochemistry for phosphorylated histone H3 (pH3) in intestine sections of tumor-bearing fish and the sibling controls at 5 dpf. pH3 (K, L), DAPI (M, N) and EGFP (O, P) images are shown. White arrows indicate intestinal cells positive for pH3, EGFP, and DAPI. In the merged images (Q, R), pH3, DAPI and EGFP signals are shown in red, blue and green, respectively. Scale bar indicates 100 μm. (S) The number of intestinal cells positive for pH3, EGFP, and DAPI. The number of pH3, EGFP, and DAPI positive cells was counted from single section per individual fish. The data harbors 8 and 6 biological replicates from tumor-bearing fish and the sibling controls, respectively. Error bars represent ± s.e.m.

Upon closer examination of *kras*^*G12D*^-expressing posterior intestine, we found that *pInt-Gal4* was expressed in cdh1 (E-cadherin)-positive intestinal cells (Fig. 3A-3H), indicating that expression of *pInt-Gal4* occurred specifically in epithelial cells in the posterior intestine. Fig. 3A-3H demonstrated that intestinal epithelial cells outgrew apically while the basal membrane structure seemed unaffected with hematoxylin and eosin (HE) staining supporting these findings (Fig. 3I-3L). Based on these atypia phenotypes, it was likely that *pInt-Gal4*-driven *kras*^*G12D*^ expression in the posterior intestine led to dysplasia, a type of tumor. Despite the disorganized structure of posterior intestine, the intestinal lumen was not completely disrupted (Fig. 3I-3L). Consistent with this, food was present in the intestinal lumen of tumor-bearing fish following feeding (Fig. S3A-S3B).

**Figure 3.**
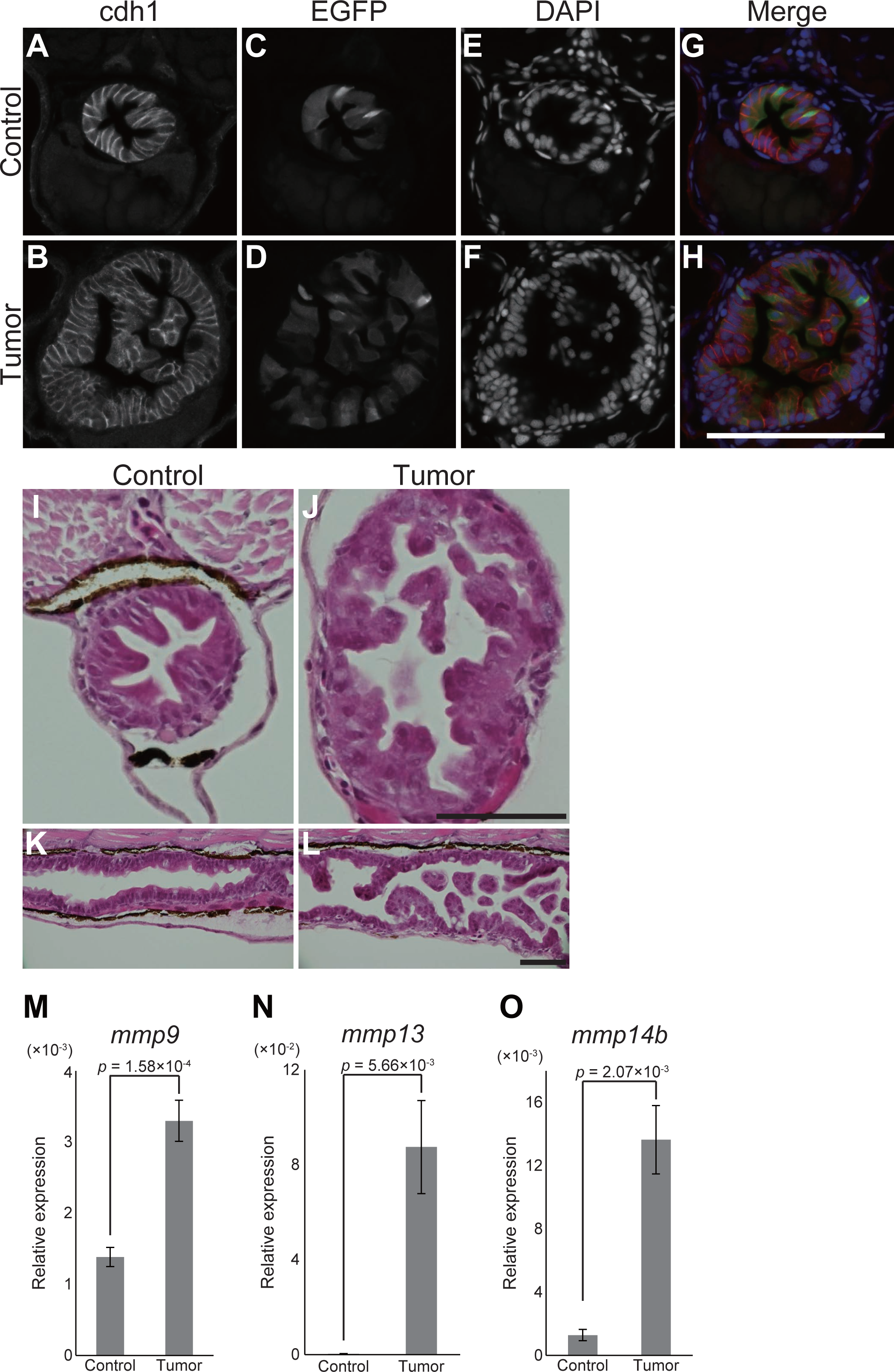
*pInt-Gal4* driven *kras*^*G12D*^ expression results in dysplasia originating from intestinal epithelial cells. (A)-(H) Representative images of fluorescent immunohistochemistry for cdh1 in intestine sections of the sibling controls and tumor-bearing fish at 5 dpf. cdh1 (A, B), EGFP (C, D) and DAPI (E, F) images are shown. In the merged images (G, H), cdh1, EGFP and DAPI signals are shown in red, green and blue, respectively. Scale bar indicates 100 μm. (I)-(L) Representative images of HE-stained intestine sections of the sibling controls (I, K) and tumor-bearing fish (J, L) at 5 dpf. Transversal and sagittal sections are shown in (I, J) and (K, L), respectively. Scale bar represents 50 μm. (M)-(O) qPCR analysis for *mmp* genes in the intestine at 9 dpf. The scores are normalized to expression of *rpl13a*. The data harbors 5 biological replicates, each containing the intestines from 5 fish. Error bars represent ± s.e.m. Statistical significance was tested using student’s *t*-test (unpaired, one-tailed).

We did not observe visible invasion and dissemination of EGFP positive cells in our experimental window (Fig. 3A-3L). Despite this, qPCR experiments demonstrated that expression of matrix metalloproteinases genes (*mmp9*, *mmp13* and *mmp14b*) was strongly increased in *kras*^*G12D*^-expressing intestinal cells, a molecular clue for invasiveness of tumor cells (Fig. 3M-3O) (Hanahan and Weinberg, 2011). Altogether, these suggest that the detected outgrowth of intestinal epithelial cells resulted in formation of dysplasia, and thus an intestinal tumor. According to the histological definitions for malignant tumor (cancer), lack of invasion and metastasis implicate that the intestinal tumor might be benign. However, because our following analyses revealed systemic adverse effects on host by the intestinal tumor, we in this manuscript simply define our model as an intestinal tumor model. Collectively, we found a combination of the *Gal4* line and oncogene that drives the intestinal tumor at an early stage of zebrafish development.

### Zebrafish intestinal tumor causes local and distant inflammation

In addition to the classical definitions for cancer (malignant tumor), recent advances in molecular biology have revealed a set of molecular features that is useful to characterize cancer, known as the hallmarks of cancer (Hanahan and Weinberg, 2011). For example, it is known that cancer recruits innate immune cells such as neutrophils for survival and for promoting metastasis, and that cancer causes systemic, distant inflammation, phenomena observed across species including human patients (Fearon et al., 2012; Hanahan and Weinberg, 2011; McAllister and Weinberg, 2014). Importantly, zebrafish models have played important roles in this field, providing significant insights into the dynamics of innate immune cells in such as tumor initiation in vivo (Feng et al., 2012; Feng et al., 2010; Mione and Zon, 2012; Patton, 2012). In order to determine if the intestinal tumor recruits neutrophils and causes systemic inflammation, we generated tumor-bearing fish carrying *Tg(lyz:EGFP)*, which expresses EGFP in neutrophils (Kitaguchi et al., 2009).

Microscopic analyses showed considerable increase for the number neutrophils at the whole-organismal level in tumor-bearing fish at 7 dpf (Fig. 4A-4H). Immunostaining with anti-Lyz antibody revealed that neutrophils were accumulated in the intestinal tumor when compared to the normal intestine (Fig. 4I-4O). During the cause of the experiments, we noted that neutrophils had also infiltrated the liver (Fig. 4P-4Q). In order to better visualize tumor-induced liver inflammation, mCherry was expressed specifically in the liver using the liver specific *fabp10a* promoter (*Tg(fabp10a:mCherry)*) (Fig. 4P-4Q) (Her et al., 2003). We counted the number of EGFP-positive neutrophils in the liver expressing mCherry. As a result,we found that the number of neutrophils in the intestines of tumor-bearing fish was greater than that in the sibling controls (12 ± 2.3 vs 30 ± 6.0, *p* = 0.0062: Fig. 4P-4Q). With respect to local and systemic inflammation, the intestinal tumor we developed appeared to harbor a feature of cancer (malignant tumor). Furthermore, the livers of tumor-bearing fish were larger than those of their sibling controls, a phenomenon known as hepatomegaly (0.028 ± 0.0013 mm^2^ vs 0.038 ±.0.0016 mm^2^, *p* = 0.00016: Fig. 4S). Tumor-induced hepatomegaly is seen also in mammalian tumor models including a colon cancer model (Bonetto et al., 2016; Hojo et al., 2017), and human cancer patients (Lieffers et al., 2009). These results suggest that the intestinal tumor adversely affects the liver, and that the model is able to recapitulate tumor-induced phenotypes observed in mammals and human patients.

**Figure 4.**
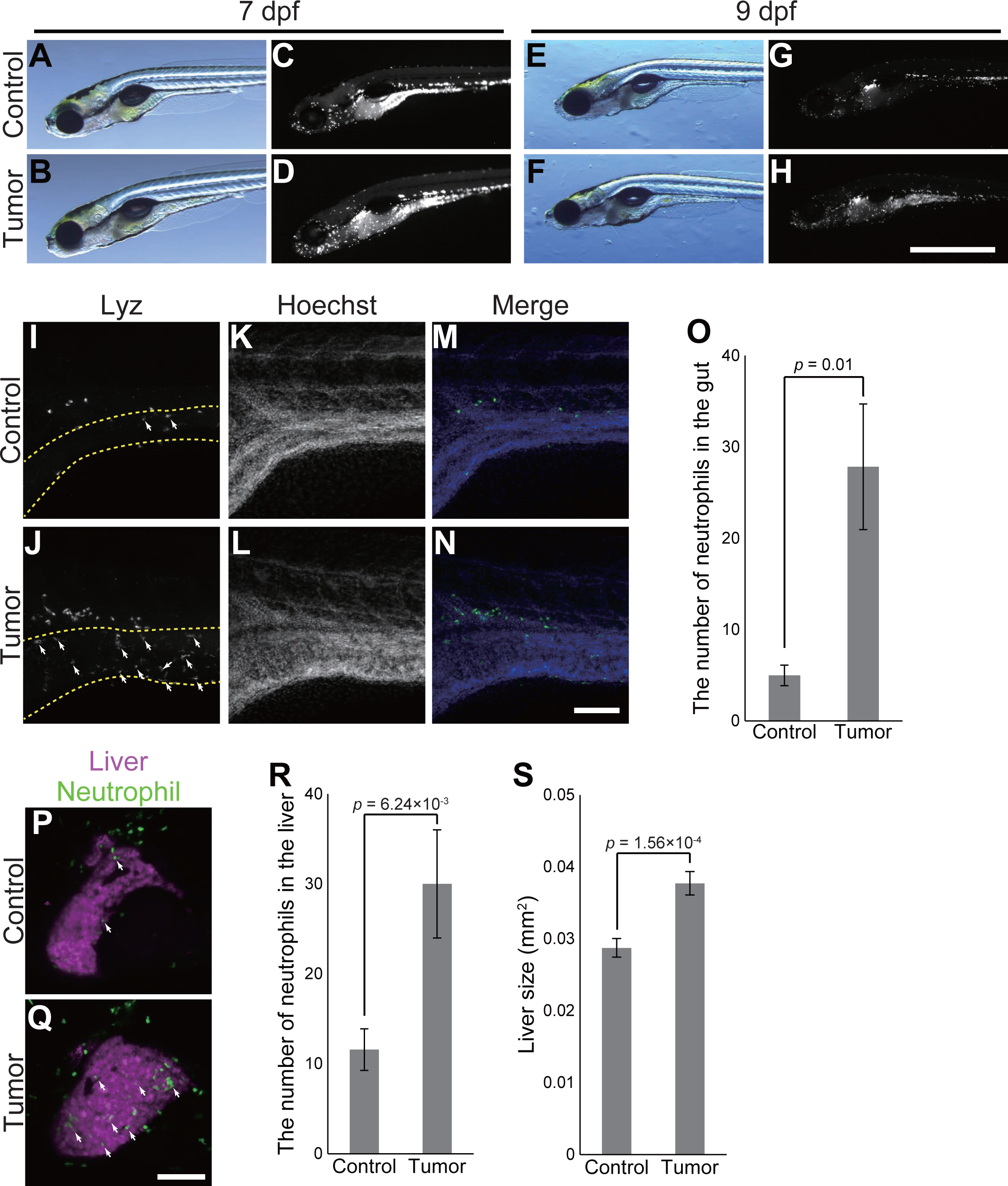
The zebrafish intestinal tumor instigates local and distant inflammation. (A)-(H) Representative images of the sibling controls and tumor-bearing fish carrying *Tg(lyz:EGFP)* transgene at 7 and 9 dpf. Bright field (A, B, E, F) and EGFP (C, D, G, H) images are shown. Scale bar indicates 1 mm. (I)-(N) Representative images of whole mount fluorescent immunohistochemistry for lyz in the intestines of the sibling controls and tumor-bearing fish at 7 dpf. Lyz (I, J) and Hoechst33342 (K, L) images are shown. The intestine is shown by yellow-dotted lines in (I, J). In the merged images (M, N), Lyz and Hoechst33324 signals are shown in green and blue, respectively. White arrows indicate representative neutrophils in the intestine. Scale bar indicates 100 μm. (O) The number of neutrophils in the intestines of the sibling controls and tumor-bearing fish.The data harbors 6 biological replicates. Error bars represent ± s.e.m. Statistical significance was tested using student’s *t*-test (unpaired, one-tailed). (P-Q) Representative images of the livers of the sibling controls and tumor-bearing fish carrying *Tg(lyz:EGFP)* and *Tg(fabp10a:mCherry)* at 7 dpf. Neutrophils and the liver are shown by green and magenta, respectively. White arrows indicate representative neutrophils in the liver. Scale bar indicates 100 μm. (R) The number of neutrophils in the livers of the sibling controls and tumor-bearing fish at 7 dpf. The data harbors 12 biological replicates. Error bars represent ± s.e.m. Statistical significance was tested using student’s *t*-test (unpaired, one-tailed). (S)Liver size of the sibling controls and tumor-bearing fish at 7 dpf. Liver size was measured from *Tg(fabp10a:mCherry)* images using ImageJ software. The data harbors 12 biological replicates. Error bars represent ± s.e.m. Statistical significance was tested using student’s *t*-test (unpaired, one-tailed).

### Zebrafish intestinal tumor impedes organismal growth and causes organismal death

Next, to further demonstrate utility of the novel intestinal tumor model, we aimed to identify other systemic effects caused by the intestinal tumor. We found that tumor-bearing zebrafish were significantly smaller than the sibling controls (Figs. 5A and S4), the difference observable from 7 dpf. The results varied among clutches at 7 dpf, whereas the growth defect phenotype was very consistent at 9 dpf (Fig. S4A-S4B). The growth defect phenotype was identified in the complete absence of foods (i.e. exogenous nutrient): although zebrafish larvae are able to eat from 5-6 dpf, yolk-derived nutrient inherited from the mother keep fish alive without visible abnormalities at least until 9 dpf. This enabled us to ignore experimental variations on zebrafish behaviors related to eating and on nutrient absorption rate in the intestine in explaining the growth defect phenotype. Based on these analyses, we concluded that the local intestinal tumor caused a systemic growth defect.

**Figure 5.**
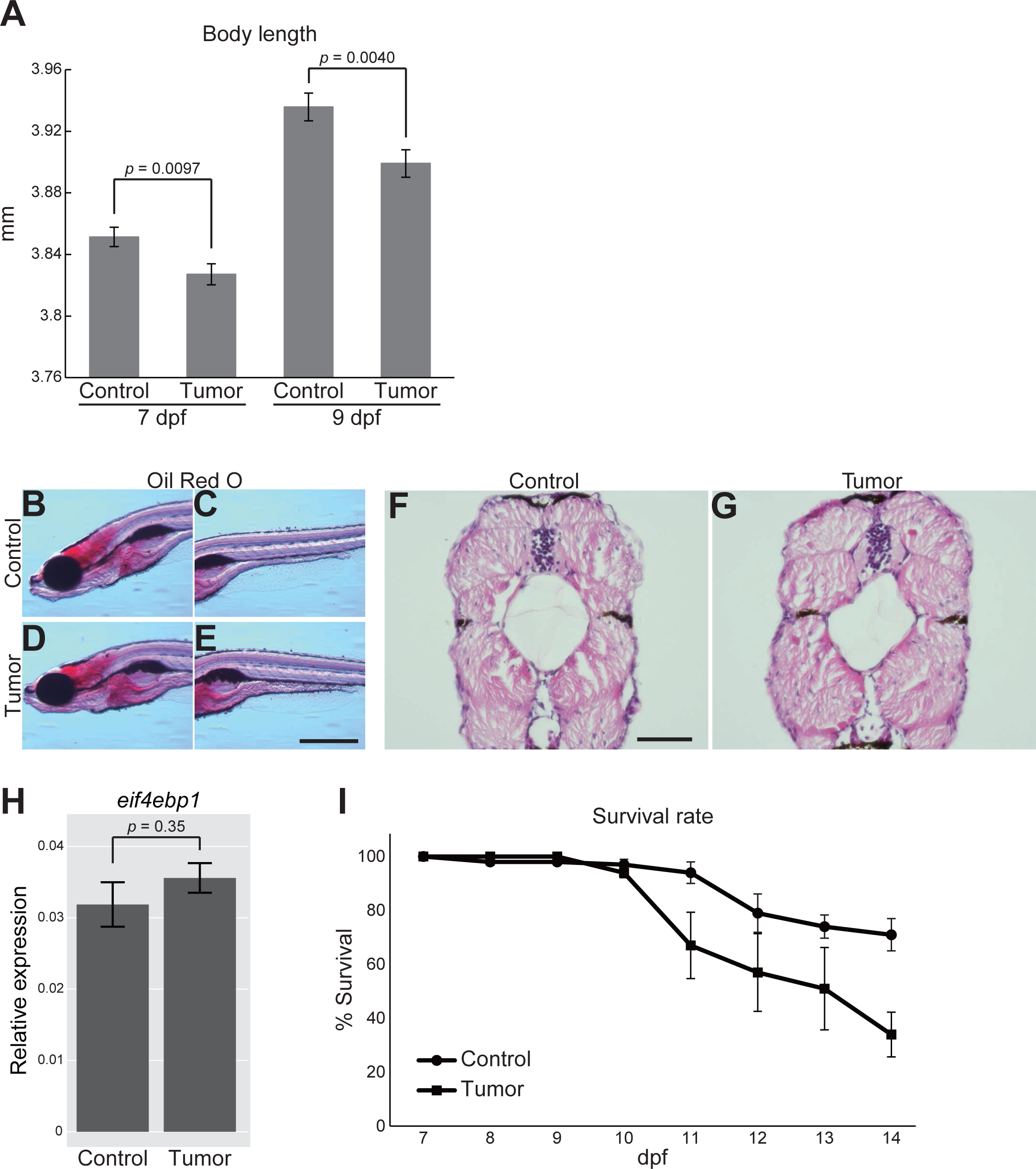
The zebrafish intestinal tumor causes the systemic growth defect and organismal death. (A) Body length data of the sibling controls and tumor-bearing fish at 7 and 9 dpf. The number of fish used is 163 (7 dpf control fish), 155 (7 dpf tumor-bearing fish), 154 (9 dpf control fish) and 154 (9 dpf tumor-bearing fish). Data from three independent clutches are pooled. Data from each clutch are shown in Fig. S4. Error bars represent ± s.e.m. Statistical significance was tested using student’s *t*-test (unpaired, two-tailed). (B)-(E) Representative images of Oil Red O staining for the sibling controls (B, C) and tumor-bearing fish (D, E) at 9 dpf. Scale bar represents 500 μm. Red-stained areas represent endogenous lipids in fish. (F)-(G) Representative images of HE-stained body sections for the sibling controls (G) and tumor-bearing fish (H) at 9 dpf. Scale bar represents 50 μm. (H) qPCR analysis for *eif4ebp1* in the body (without the intestine or intestinal tumor) in the sibling controls and tumor-bearing fish at 9 dpf. Scores are normalized to expression of *rpl13a*. The data harbors 5 biological replicates, each containing 5 fish. Error bars represent ±s.e.m. Statistical significance was tested using student’s *t*-test (unpaired, two-tailed). (J) Survival analysis of the sibling controls and tumor-bearing fish. Twenty fish per a tank were fed from 7 dpf, and the number of dead fish was counted everyday. Data were obtained by five independent experiments. Error bars represent ± s.e.m.

It is well-known that tumor-bearing animals waste muscle and fat, resulting in a loss of weight (i.e. tumor-induced cachexia) (Das et al., 2011; Fearon et al., 2012; Figueroa-Clarevega and Bilder, 2015; Kwon et al., 2015). In fact, Kwon et al. find that fly tumor alters homeostasis of systemic lipids including triglyceride TG (Kwon et al., 2015). To explore whether the growth defect phenotype could be attributed to cachexia, Oil Red O staining for neutral TGs and lipids was performed. Stronger staining was detected for the liver and brain at 9 dpf, a pattern of which was not prominently different between tumor-bearing fish and the sibling controls (Fig. 5B-5E). This suggested that the intestinal tumor at this stage did not have a strong impact on the systemic lipid level. In addition, HE staining did not find obvious loss of host tissues such as muscles at 9 dpf (Fig. 5F-5G). These were consistent with qPCR data showing that *eif4ebp1*, a marker for reduced insulin signaling (Figueroa-Clarevega and Bilder, 2015; Kwon et al., 2015), was not affected by the intestinal tumor (Fig. 5H). Thus, the growth defect phenotype we identified was unlikely to be canonical cachexia (Figueroa-Clarevega and Bilder, 2015; Kwon et al., 2015).

Next we asked if the intestinal tumor worsens mortality of zebrafish. We counted the number of dead and live fish every day and found that the survival rate of tumor-bearing fish (less than 50% at 14 dpf) was significantly lower than that of the sibling controls (approximately 80%; Fig. 5I). This phenotype was not due to a complete defect in swimming ability and/or a complete loss of appetites in tumor-bearing fish, because tumor-bearing fish were able to swim and eat (Figs. S3A-S3B). Importantly, visible metastases were still not detected by microscopic inspection at 14 dpf (unpublished observation), indicating that the local intestinal tumor affected the survival rate.

Taken together, the intestinal tumor driven by strong oncogene *kras*^*G12D*^ expression was histologically classified as dysplasia, a type of benign tumor, but yet detrimental for organismal physiology, causing inflammation, hepatomegaly, growth defects, and organismal death. Practically, our novel intestinal tumor model is useful in that the major systemic phenotypes, which are clinically observed, occur within 2 weeks after fertilization, when zebrafish larvae are still small and transparent.

### Zebrafish intestinal tumor reduces hepatic cyp7a1 expression and lowers the bile acids synthesis

To examine the effects of the intestinal tumor on host at the gene expression level and identify differentially expressed genes (DEGs), whole-organismal level RNA-seq experiments were performed. Zebrafish at 7 dpf were roughly dissected into the three parts, the liver, the intestinal tumor or normal intestine, and the rest part of body (Fig. 6A and Table S2-S6). We were particularly focused on the liver since the liver was preferentially inflamed by the intestinal tumor (Fig. 4), despite a lack of visible metastasis to the liver in our experimental setting. A set of genes potentially affected by the intestinal tumor (Table S2-S6) was used for further validation by qPCR to identify consistently affected genes: RNA-seq experiments served as a screening to find candidate DEGs.

**Figure 6.**
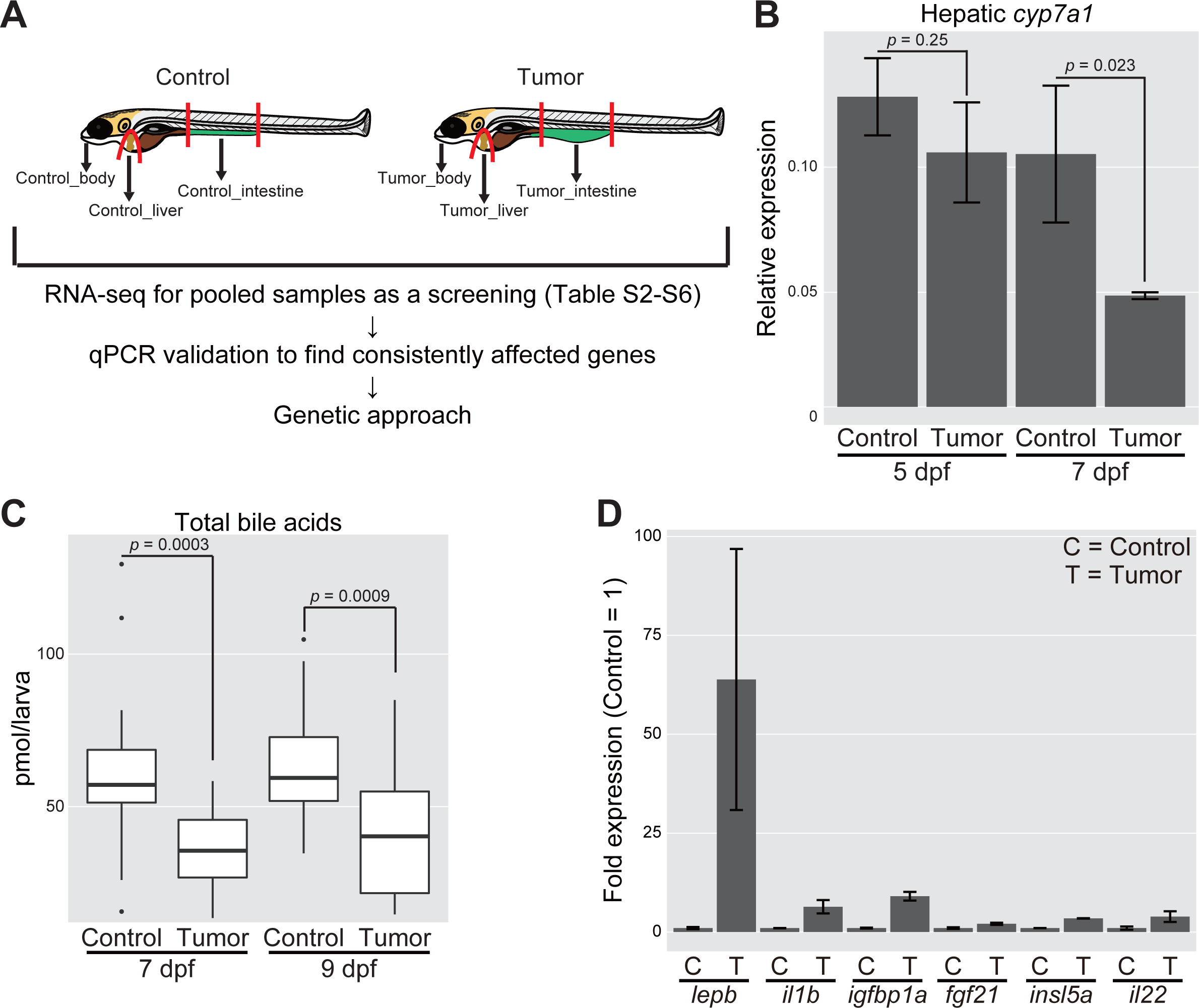
Whole-organismal level gene expression analysis identifies tumor-liver crosstalk characterized by the decreased expression of hepatic *cyp7a1* and the reduced amount of total bile acids. (A) Schematic representation of zebrafish dissection in our RNA-seq experiments followed by qPCR validation and genetics. (B) Expression of *cyp7a1* in the liver. The scores are normalized to expression of *rpl13a*. The data harbors 3 biological replicates, each containing 7 fish for 5 dpf and 5 fish for 7 dpf, respectively. Error bars represent ± s.e.m. Statistical significance was tested using student’s *t*-test (unpaired, one-tailed). (C) Measurement for systemic bile acids levels at 7 and 9 dpf. The number of fish used is 19 (7 dpf control fish), 19 (7 dpf tumor-bearing fish), 22 (9 dpf control fish) and 18 (9 dpf tumor-bearing fish). Statistical significance was tested using student’s *t*-test (unpaired, one-tailed). (D) Expression of a set of secreted protein-coding genes in the intestinal tumor and normal intestine. The scores are normalized to expression of *rpl13a* and to the sibling controls (=1). The data harbors 3 biological replicates, each containing 5-7 fish. Error bars represent ± s.e.m.

Notably, we found that hepatic *cyp7a1*, the gene encoding the rate-limiting enzyme that acts at the 1st step of converting cholesterol to bile acids (BAs) (Kuipers et al., 2014; Thomas et al., 2008), was decreased in the presence of the intestinal tumor (Fig. 6B). Total BAs were then individually quantified, and fish from multiple clutches were analyzed to test whether reduced *cyp7a1* expression resulted in a consequent drop in BAs. The colorimetric quantitative assay demonstrated that total BAs levels were significantly reduced (∼50%) in tumor-bearing fish both at 7 dpf and 9 dpf (Figs. 6C). Despite the reduction in total BAs levels, total cholesterol levels were not significantly affected by the intestinal tumor (Fig. S5A). These data suggested that the zebrafish intestinal tumor disrupts the hepatic BAs synthesis via down-regulation of *cyp7a1* in the liver, anomaly that could account for the systemic phenotypes caused by the intestinal tumor.

We next analyzed our RNA-seq data on the normal intestine and the intestinal tumor. Comparison between these two samples identified a set of genes strongly elevated in the intestinal tumor (Fig. 6D). DEGs included inflammatory response genes including *interleukin 1b* (*il1b*) and *matrix metallopeptidase 13* (*mmp13*), and *myeloid-specific peroxidase* (*mpx*) (a marker for neutrophils and macrophages), which were in line with our imaging data (Fig. 4I-O), and known RAS targets such as *gamma-glutamyltranspeptidase1* (*ggt1*) (Figs. 3N and S5B-S5C). Moreover, several secreted factors were elevated including *leptin b* (*lepb*), *insulin-like growth factor binding protein 1a* (*igfbp1a*), *insulin-like peptide 5 a/b* (*insl5a and b*), *fibroblast growth factor 21* (*fgf21*), *interleukin 22* (*il22*), and *il1b* (Fig. 6D). The secreted protein-coding genes up-regulated in the intestinal tumor were considered as promising candidates that may reduce the production of hepatic BAs and/or underlie the systemic phenotypes. Fgf19 and Fgf21 in mice have a role in controlling BAs synthesis (Degirolamo et al., 2016). Insulin antagonist ImpL2 causes cachexia in *Drosophila*, and IGFBPs have been implicated in mammalian cancers (Baxter, 2014; Figueroa-Clarevega and Bilder, 2015; Kwon et al., 2015). *insl5* encodes a peptide that belongs to a relaxin family as well as fly Dilp8 (Burnicka-Turek et al., 2012; Grosse et al., 2014). Mouse studies reported a role for Insl5 in glucose homeostasis and the orexigenic signaling, but its function in tumor-associated pathology is unknown (Burnicka-Turek et al., 2012; Grosse et al., 2014). It is also possible that inflammatory cytokines such as *il1b* and *tnf* reduces expression of hepatic *cyp7a1* (Okin and Medzhitov, 2016). Overall, the whole-animal level RNA-seq experiments and qPCR revealed the intriguing abnormality in the liver metabolism coincident with de-regulated expression of secreted protein-coding genes in the intestinal tumor.

### Driving cyp7a1 expression in the liver ameliorates tumor-induced liver inflammation

In order to ask whether the reduced *cyp7a1* expression in the liver affects tumor-induced systemic phenotypes, we generated a transgenic line expressing *cyp7a1* under the control of the *fabp10a* promoter (Her et al., 2003). Expression of *cyp7a1* was linked to mCherry with P2A (Kim et al., 2011) (Fig. 7A). The transgene expression was ascertained by microscopic observation and qPCR (Fig. 7B-7D). Overexpression of *cyp7a1* in the liver significantly restored total BAs levels both at 7 and 9 dpf in tumor-bearing fish (Figs. 7E-7F). The transgene also tended to increase total BAs in their tumor-free sibling controls. Altogether, the *fabp10a:mCherry-P2A-cyp7a1* transgene was able to restore the BAs production in tumor-bearing fish, further supporting that the intestinal tumor affects cholesterol-BAs flux via down-regulation of *cyp7a1*.

**Figure 7.**
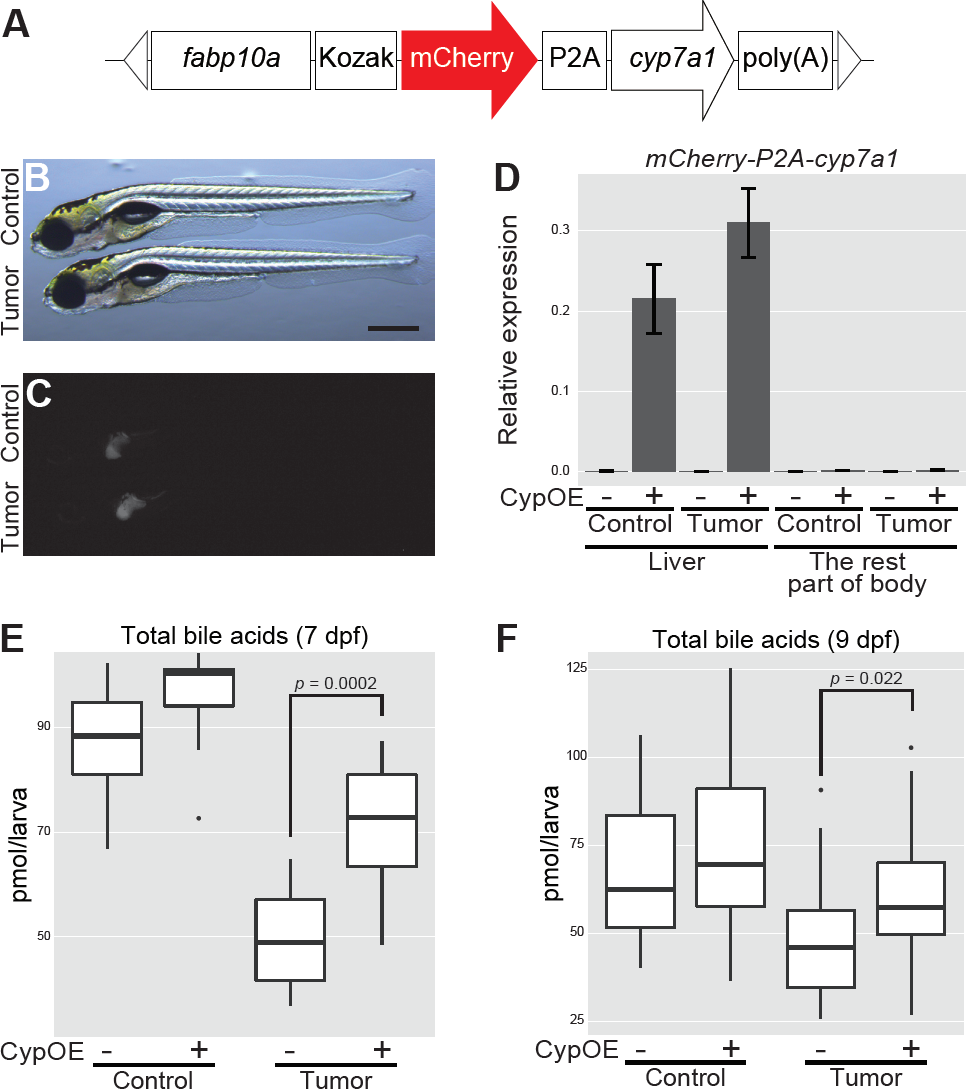
Overexpression of *cyp7a1* in the liver restores the amount of total bile acids in tumor-bearing fish. (A) The structure of *fabp10a:mCherry-P2A-cyp7a1*. The white triangles represent the recognition sequence by I-SceI meganucleases. (B)-(C) Representative images of the *mCherry-cyp7a1* transgene expression in the liver. Control refers to *Tg(pInt-Gal4)*^*+/Tg*^; *Tg(UAS:EGFP)*^*+/Tg*^; *Tg(fabp10a:mCherry-P2A-cyp7a1)*^*+/Tg*^ while tumor-bearing fish to *Tg(pInt-Gal4)*^*+/Tg*^; *Tg(5×UAS:EGFP-P2A-kras*^*G12D*^)^*+/Tg*^; *Tg(fabp10a:mCherry-P2A-cyp7a1)*^*+/Tg*^. Scale bar represents 500 μm. Bright field (B) and mCherry (C) images are shown. (D) qPCR analysis for detecting *mCherry-cyp7a1* mRNAs in the liver and the rest part of the body at 7 dpf. The scores are normalized to expression of *rpl13a*. The data harbors 3 biological replicates, each containing 3 fish. Error bars represent ± s.e.m. CypOE - and + indicate the absence and presence of *Tg(fabp10a:mCherry-P2A-cyp7a1)*, respectively. (E)-(F) Measurement for systemic bile acids levels at 7 (n = 10 per a group) and 9 dpf (n = 30-31 per a group). Statistical significance was tested using student’s *t*-test (unpaired, one-tailed). CypOE - and + indicate the absence and presence of *Tg(fabp10a:mCherry-P2A-cyp7a1)*, respectively.

These results promoted us to test if overexpression of *cyp7a1* in the liver could rescue the intestinal tumor-induced systemic phenotypes. We examined whether three major tumor-induced phenotypes, liver inflammation, hepatomegaly, and the growth defect, were rescued by the *fabp10a:mCherry-P2A-cyp7a1* transgene (Fig. 8). We found that *cyp7a1* overexpression did not significantly rescue the growth defect phenotype (Fig. 8A). As was the case for Fig. 5A, the results to some extent varied depending on clutches: in one clutch, we observed a trend for the rescue while not in a different clutch. Upon pooling data from multiple clutches, we concluded that *cyp7a1* overexpression did not consistently and significantly rescue the growth defect phenotype. Moreover, tumor-induced hepatomegaly (0.028 ± 0.0013 mm^2^ (control) vs 0.038 ± 0.0016 mm^2^ (tumor), *p* = 0.00016: Fig. 4S) was not affected by *cyp7a1* overexpression in the liver (0.028 ± 0.0011 mm^2^ (control) vs 0.033 ± 0.0018 mm^2^ (tumor), *p* = 0.012: Fig. 8B-8D).

**Figure 8.**
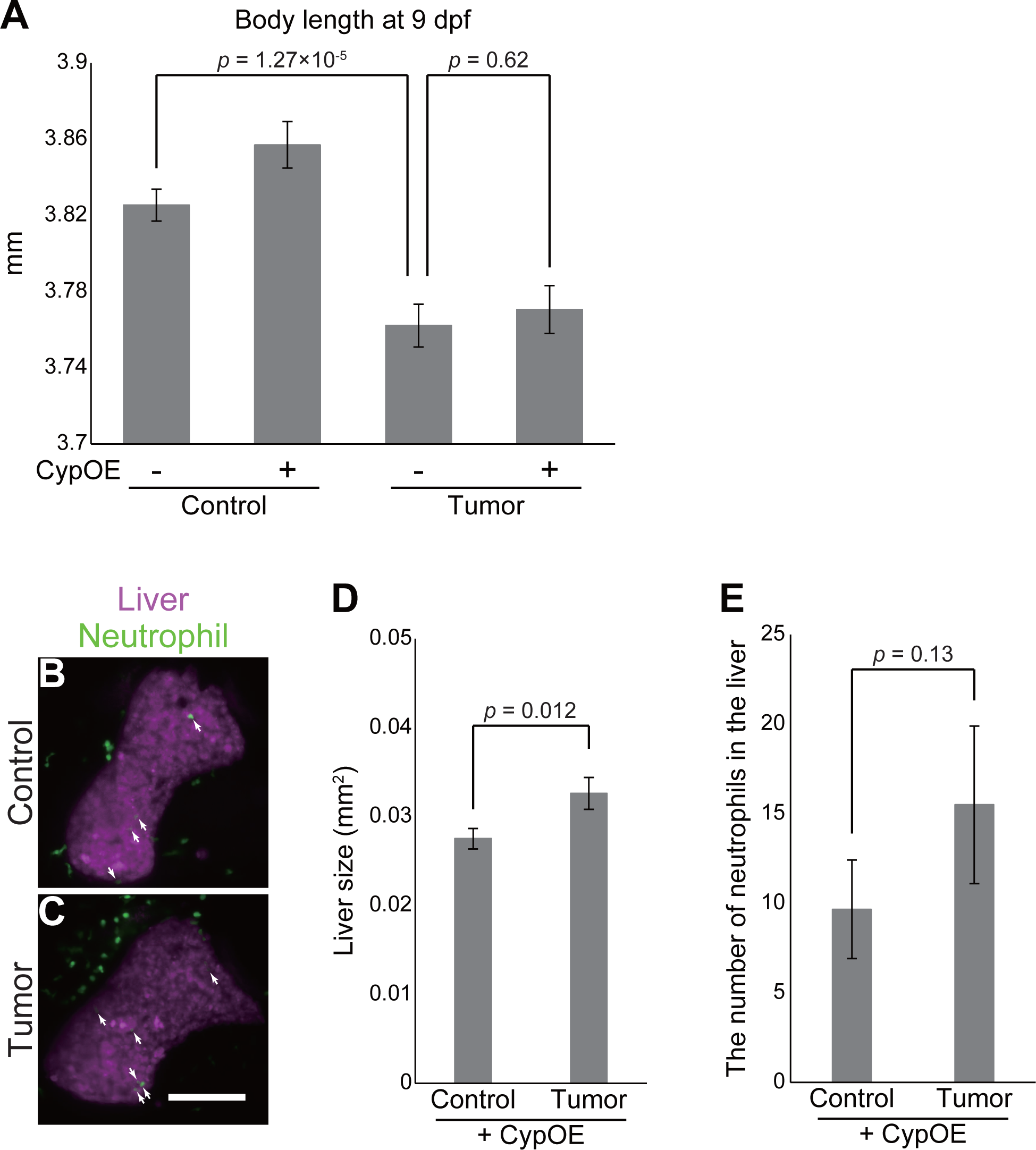
Overexpression of *cyp7a1* in the liver ameliorates tumor-induced liver inflammation. (A) Body length data of the sibling controls and tumor-bearing fish at 9 dpf in the *Tg(fabp10a:mCherry-P2A-cyp7a1)* background. The number of fish used is 79 (control fish), 73 (control fish with *Tg(fabp10a:mCherry-P2A-cyp7a1*)), 81 (tumor-bearing fish) and 74 (tumor-bearing fish with *Tg(fabp10a:mCherry-P2A-cyp7a1*)). Error bars represent ± s.e.m. Statistical significance was tested using student’s *t*-test (unpaired, two-tailed). CypOE - and + indicate the absence and presence of *Tg(fabp10a:mCherry-P2A-cyp7a1)*, respectively. (B)-(C) Representative images of the livers of the sibling controls and tumor-bearing fish carrying *Tg(lyz:EGFP)* and *Tg(fabp10a:mCherry-P2A-cyp7a1)* at 7 dpf. Neutrophils and the liver are shown by green and magenta, respectively. Scale bar indicates 100 μm. (D) Liver size and (E) the number of neutrophils of the sibling controls and tumor-bearing fish carrying *Tg(lyz:EGFP)* and *Tg(fabp10a:mCherry-P2A-cyp7a1)* at 7 dpf. Liver size was measured from *Tg(fabp10a:mCherry-P2A-cyp7a1)* images using ImageJ software. The data harbors 18 biological replicates. Error bars represent ± s.e.m. Statistical significance was tested using student’s *t*-test (unpaired, one-tailed).

Interestingly, the number of neutrophils observed in the liver was comparable between the sibling controls and tumor-bearing fish in the *Tg(fabp10a:mCherry-P2A-cyp7a1)* background (9.7 ± 2.8 (control) vs 16 ± 4.4 (tumor), *p* = 0.134: Fig. 8B-8C, and 8E) in contrast to our data in the *Tg(fabp10a:mCherry)* background (12 ± 2.3 (control) vs 30 ± 6.0 (tumor), *p* = 0.0062: Fig. 4P-4Q). As an important detail, these experiments (Figs. 4R-4S and 8B-8E) were performed using staged-matched fish (7 dpf), which was demonstrated by that liver size and the number of neutrophils were similar in the control groups. Despite statistically insignificant, there was still a trend for the increase in the number of neutrophils in tumor-bearing fish in the *Tg(fabp10a:mCherry-P2A-cyp7a1)* background. This might suggest that the rescue by *Tg(fabp10a:mCherry-P2A-cyp7a1)* was partial, consistent with that the extent of rescue for total BAs levels were not 100 % (Fig. 7E-7F). Alternatively, another factor might contribute to liver inflammation by the intestinal tumor.

*cyp7a1* has not been considered as a crucial host gene in tumor-induced distant inflammation. Yet, studies in different contexts support our observation that the intestinal tumor actively reduces expression of hepatic *cyp7a1* to promote liver inflammation (Fig. 9). In mice, overexpression of Cyp7a1 in the liver suppresses lipopolysaccharide (LPS)-induced hepatic inflammation and fibrosis (Liu et al., 2016). It is also known that sustained inflammation reduces expression of Cyp7a1, suggestive of a role for Cyp7a1 in inflammation in mice (Okin and Medzhitov, 2016). Collectively, the current study, as the demonstration for utility of the model, identifies *cyp7a1* as a host gene that mediates liver inflammation, one of tumor’s adverse effects on host by the intestinal tumor.

**Figure 9.**
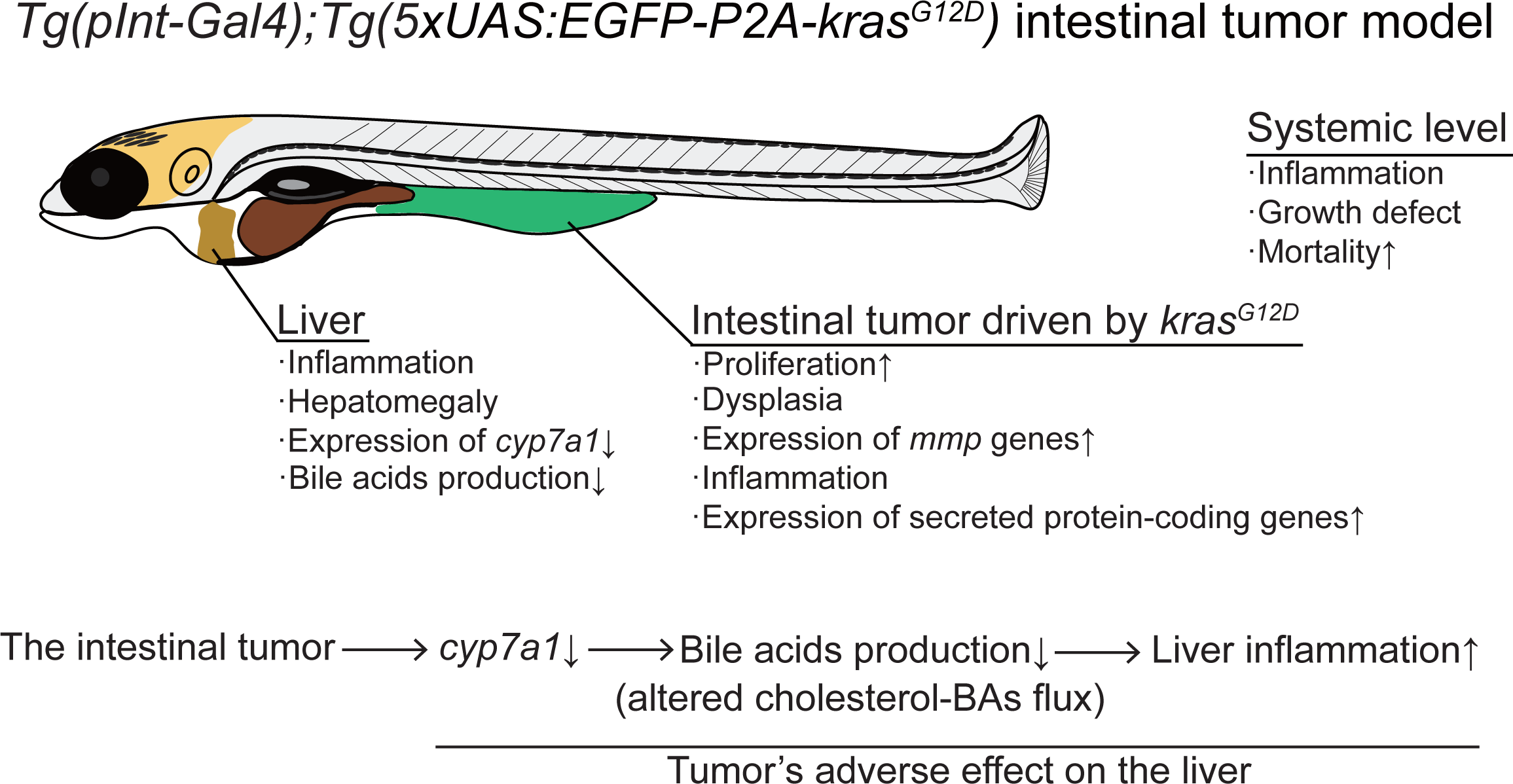
The graphical summary of this study. *kras*^*G12D*^ expression driven by *pInt-Gal4* results in dysplasia in the posterior intestine. Despite histologically benign and restricted to the intestine, the intestinal tumor causes a set of systemic adverse effects on host. The intestinal tumor recruits neutrophils to the liver, accompanied with hepatomegaly. Tumor-bearing fish grow less than the sibling controls do, and die around at 14 dpf. The intestinal tumor communicates with the liver, altering cholesterol-BAs flux. This interaction is important for tumor-induced liver inflammation, but not for other phenotypes.

## Discussion

This study has two major advances. First, we established the novel zebrafish intestinal tumor model, which is optimized for studying body-wide tumor-organ interaction in vivo. Second, using the model, we discovered a tumor-liver interaction that mediates enhanced recruitment of neutrophils to the liver in tumor-bearing fish, via down-regulation of a cholesterol-metabolizing gene *cyp7a1* as a critical host gene.

### Establishment of a novel intestinal tumor model in zebrafish

The zebrafish intestinal tumor model we have newly established harbors several strengths for studying tumor-organ interaction at the whole-organismal level (Fig. 7). The combination of *pInt-Gal4* and UAS-controlled *kras*^*G12D*^ induces epithelial tumor formation in the posterior intestine at as early as 5 dpf, when zebrafish are small and completely transparent (Figs. 1-3). Yet, zebrafish larvae after 5 dpf are able to swim and eat and therefore it is likely that essential organs such as the liver are already mature at this time point. Even though the intestinal tumor is histologically dysplasia (not fully malignant), the intestinal tumor causes detrimental effects on host including systemic inflammation, hepatomegaly, a growth defect, metabolic defects, and organismal death (Figs. 4-8). The model even made it possible to visualize the intestinal tumor-induced inflammation in the liver of live fish (Fig. 4). Furthermore, the growth defect phenotype we discovered does not depend on exogenous food intake, simplifying our investigation on how the intestinal tumor causes systemic growth defect (Fig. 5).

To date, a genetically engineered, robust zebrafish intestinal tumor model has not yet been available (Lobert et al., 2016). The structure of intestinal tract in zebrafish is different from mice and humans, especially in that zebrafish lacks the stomach. Still, the zebrafish intestine shares common features with mammalian intestines, the notion validated by anatomical analysis and comprehensive gene expression study (Lobert et al., 2016; Wallace et al., 2005; Wang et al., 2010). On the basis of these, the zebrafish intestine appears to be analogous to the small intestine, colon, and rectum of mammals. Relevance to human diseases of our model is also supported by that the intestinal tumor model exhibits liver phenotypes observed in murine colon tumor models such as *Apc*^*Min/+*^ and human patients (Bonetto et al., 2016; Lieffers et al., 2009; Narsale et al., 2015): it is of note that *Apc*^*Min/+*^ is a model of adenoma (histologically benign) and potent to cause adverse effects on host. Therefore, histological classification of tumors (benign or malignant) does not always correlate with the degree of adverse effects on host. Taken together, we expect that our model will be a valuable tool for studying biology of intestinal tumors.

It is also important to note that there are other zebrafish models that develop tumors at an early stage of zebrafish development, which are thus potentially useful for studying tumor-organ crosstalk at the whole organismal level. For instance, Mione and colleagues established a novel brain tumor model using *HRAS*^*V12*^, in which increased brain size was observed already at 3 dpf (Mayrhofer et al., 2017). Activating beta-catenin signal promotes liver enlargement associated with enhanced proliferation at 6 dpf in the model established by Stainier and colleagues (Evason et al., 2015). These models are definitely useful to obtain insights into how various types of local tumors affect developing vertebrates.

### Identification of a tumor-induced growth defect in developing zebrafish

Our model exhibits an intriguing systemic phenotype: tumor-bearing fish do not grow well compared to their sibling controls (Fig. 5A). This phenotype was neither accompanied with a clear reduction of the systemic lipid level (Fig. 5B-5E) nor with reduced insulin signaling (Fig. 5H), common phenotypes observed in cachexia patients and animal models (Fearon et al., 2012; Figueroa-Clarevega and Bilder, 2015; Kwon et al., 2015). Hence, we at this point assume that the observed growth defect is not the typical tumor-induced cachexia.

The growth defect phenotype to some extent resembled the growth delay in flies harboring an imaginal disc tumor or local wounds (Colombani et al., 2015; Colombani et al., 2012; Garelli et al., 2012; Garelli et al., 2015; Katsuyama et al., 2015; Owusu-Ansah and Perrimon, 2015; Vallejo et al., 2015). Secreted fly-specific peptide Dilp8 and its receptor Lgr3 are at the core of adaptation of growth and developmental timing to local disruptions. Dilp8 interacts with Lgr3 expressed in neurons that are projected to the prothoracic gland to control biosynthesis of ecdysone, one of the master regulators for fly development (Colombani et al., 2015; Garelli et al., 2015; Vallejo et al., 2015). However, whether similar growth retardation occurs in vertebrate tumor models has not been validated. Our study demonstrates the first vertebrate model in which the local intestinal tumor impedes organismal growth. Secreted protein-coding genes such as *insl5a* up-regulated in the intestinal tumor may act as an upstream of the growth defect (Fig. 6D-6E).

Recent advances in pediatric oncology have greatly improved the survival rate of childhood cancer patients. Importantly, it is known that survivors of childhood cancers often have “late complications,” long-lasting (sometimes for 40-years) complications including growth defects (Robison and Hudson, 2014; Rose et al., 2016). Cancers by themselves and/or cancer treatments (e.g. chemotherapy) may cause late complications, but the details are still unknown. Our model develops the intestinal tumor at a juvenile stage when zebrafish larvae grow massively. The study thus points out the possibility that local tumor could be a cause for long-lasting growth defects in human cancer patients. This can be directly addressed once we have the ability to cure the intestinal tumor in our model so that we can test if the growth defect lasts even after removal of the intestinal tumor.

### The intestinal tumor remotely alters systemic cholesterol-BAs homeostasis through cyp7a1-mediated tumor-liver interaction to promote liver inflammation

One of the strengths of our model is that the intestinal tumor causes systemic effects when zebrafish larvae are small enough for the whole-body analysis (Figs. 1-2). This enabled us to perform whole-organismal transcriptome analysis to capture gene expression changes in the intestinal tumor and the remaining normal organs (Fig. 6 and Tables S2-S6). We found that the liver responded to the intestinal tumor in the most sensitive manner in our model (Figs. 4 and 6). In addition to tumor-induced systemic inflammation and hepatomegaly (Fig. 4) (Egeblad et al., 2010; Fearon et al., 2012; McAllister and Weinberg, 2014), hepatic expression of *cyp7a1*, the gene encoding the rate-limiting enzyme for synthesizing bile acids (BAs) (Kuipers et al., 2014; Thomas et al., 2008), was decreased at as early as 5-7 dpf in tumor-bearing fish (Figs. 6C and 6D). This reduction was concordant with the reduced total BAs levels (Fig. 6E), which was not due to the decreased body size, as we did not find any correlation between body length and the amount of bile acids in each individual (Fig. S6). Indeed, rescuing *cyp7a1* expression in the liver by means of the *fabp10a* promoter significantly restored total BAs levels in tumor-bearing fish (Fig. 7). Intriguingly, this rescue was associated specifically with buffered liver inflammation (Fig. 8): the number of neutrophils in the liver was increased in the presence of the intestinal tumor (Fig 4P-R), which was significantly ameliorated by overexpression of *cyp7a1* in the liver (Fig. 8E-8G). These results indicate that the intestinal tumor instigates liver inflammation through modulating expression of *cyp7a1* and cholesterol-BAs flux in the liver. Given that *Tg(fabp10a:mCherry-P2A-cyp7a1)* did not rescue hepatomegaly and the growth defect, it is likely that liver inflammation is independent of these phenotypes (Figs. 8-9). Our results were in line with recent studies showing a role for murine Cyp7a1 in liver inflammation in non-cancer disease models (Liu et al., 2016; Okin and Medzhitov, 2016), indicative of a generalizable role for *cyp7a1*-mediated cholesterol-BAs metabolism in diseases. These also solidify the general utility of our novel tumor model. We emphasize that our findings are of significance in that we re-defined *cyp7a1* as a host gene critical for mediating the tumor-liver-neutrophil crosstalk in vivo. It still remains unclear whether total BAs levels and/or altered cholesterol flux affect the liver, and for what the intestinal tumor causes distant inflammation in the liver. Further extensive genetic studies are ongoing to reveal physiological significance of the altered cholesterol-BAs homeostasis in tumor-bearing fish.

### Genetics on physiological interaction between tumor and normal organ(s)

Here we provide evidence for the utility of our model by showing that *cyp7a1*-mediated tumor-liver interaction underlies altered neutrophil dynamics in the livers of tumor-bearing fish. An importance of hepatic *cyp7a1* in tumor’s adverse effects on host has not been previously appreciated. Thus, the study shows that our approach is powerful to uncover previously unknown contribution of ordinary genes in tumor-induced systemic phenotypes. Three major questions are to be solved: which tumor-derived factor(s) causes hepatic *cyp7a1* down-regulation and liver inflammation? Does *cyp7a1*-mediated liver inflammation benefits the intestinal tumor? What are other host genes responsible for systemic tumor’s adverse effects on host in this model? We are addressing these questions by combining transcriptome and genetic experiments. Further genetic dissection on such physiologically important tumor-organ interaction will help to discover a therapy(s) that ameliorates host physiology harmed by tumors.

## Materials & Methods

### Zebrafish

All animal protocols were approved by the Animal Care and Use committee of Advanced Telecommunications Research Institute International. AB line was used as the standard line. Adult fish were reared at 28°C under 14 h/10 h light/dark cycle and fed hatched brine shrimp and the Hikari Lab 130 food (KYORIN). Fish were fed twice a day except weekends and holidays (once a day). Embryos were obtained by mating male fish with female fish in a water tank and were maintained at 28°C in egg water (3% sea salts, 6.4 nM methylene blue) in a plastic petri dish. Tricaine methanesulfonate (MS-222) was used as an anesthetic reagent at the concentration of 0.008% in egg water.

### Transgenic lines and plasmid construction

The transgenic zebrafish lines, gSAIzGFFD1105A (*pInt-Gal4*), gSAIzGFFM103B (*aInt-Gal4*), gSAIzGFFD886A (*Liver-Gal4*), and gSAGFF138A (*Brain-Gal4*), were generated by Tol2-transposon mediated gene trap and enhancer trap methods as described previously (Asakawa and Kawakami, 2008; Kawakami et al., 2016). *Tg(lyz:EGFP)* were obtained from National Bioresource Project Zebrafish Core Institution under the approval of the developer (Kitaguchi et al., 2009). The constructs for generating *Tg(5×UAS:EGFP-P2A-kras ^G12D^)*, *Tg(fabp10a:mCherry)* and *Tg(fabp10a:mCherry-P2A-cyp7a1)* were generated by PCR, combining the synthesized oligos and fragments amplified from the WT genome (Her et al., 2003; Omae et al., 2013). The sequences are provided in Table S1. Generation of *Tg(5×UAS:EGFP-P2A-kras*^*G12D*^) was performed as described previously (Kawakami, 2004; Thermes et al., 2002). I-SceI meganuclease was purchased from New England Biolabs and used for generating *Tg(fabp10a:mCherry)* and *Tg(fabp10a:mCherry-P2A-cyp7a1)* (Thermes et al., 2002). The existence of mCherry or EGFP-encoding transgene was inspected using Leica M165 FC fluorescent stereoscopic microscope (Leica)

### Screening of transgenic Gal4 lines that can drive tumorigenesis

*Tg(5×UAS:EGFP-P2A-kras*^*G12D*^) line was mated to each *Gal4* line carrying heterozygous *Gal4* transgene. As an example, *Tg(5×UAS:EGFP-P2A-kras*^*G12D*^)^*+/Tg*^ fish was crossed with *Tg*(*pInt-Gal4)*^*+/Tg*^*; Tg(UAS:EGFP)*^*+/Tg*^ fish to obtain *Tg(5×UAS:EGFP-P2A-kras*^*G12D*^)^*+/Tg*^*; Tg(pInt-Gal4)*^*+/Tg*^ embryos. Expression of *kras*^*G12D*^ in the siblings was examined by EGFP expression using Leica M165 FC fluorescent stereoscopic microscope (Leica). When fish with a potentially tumorous phenotype were identified, fish with no EGFP expression from the same clutch (i.e. a clutch includes siblings born on the same day from the same parents) were considered as their sibling controls. In cases where no observable phenotype could be discerned, *kras*^*G12D*^-expressing fish and the sibling controls were discriminated based on genotyping experiments. In both cases, fish harboring either *Gal4* or *Tg(5×UAS:EGFP-P2A-kras*^*G12D*^), or none of both, served as the sibling controls. For genotyping, genomic DNA was isolated from single larva by proteinase K (Takara, 1:100 dilution) in 10 mM Tris-HCl (pH 8.0) and 50 mM KCl and used as a PCR template. Each transgene was amplified using KAPA 2G Fast HS (NIPPON Genetics). *tp53* genomic region was used as the PCR control. The primers used are listed in Table S1.

### RNA isolation, cDNA synthesis, and quantitative PCR (qPCR)

For gene expression experiments, we often pooled multiple fish in a single tube. This was for obtaining sufficient amount of high-quality RNAs especially when dissection was performed, and for lowering the risk to select outliers from the clutch. Given that single female generally produces more than 50 embryos, selecting e.g. 3 ∼ 5 fish from a clutch may give rise to unwanted bias in sample collection. Pooling multiple fish, 3 ∼ 10, depending on the size of clutches, in a single tube, and treat it as one biological replicate, is useful to reduce these risks. Total RNA was isolated using TRIzol (Thermo Fisher SCIENTIFIC) or RNeasy Mini Kit (QIAGEN). cDNA was synthesized using SuperScript III First-Strand Synthesis System (Thermo Fisher SCIENTIFIC) or Transcriptor First Strand cDNA Synthesis Kit (Roche). The obtained cDNAs were 5- or 10-fold-diluted and subjected into qPCR experiments by using LightCycler480 Instrument II system and SYBR Green Master Mix (Roche). The obtained data were analyzed using the delta-Ct method. The primers used are listed in Table S1.

### Cryosectioning and fluorescent immunohistochemistry

*Tg(5×UAS:EGFP-P2A-kras*^*G12D*^)^*+/Tg*^*; Tg(pInt-Gal4)+/Tg* and *Tg*(*pInt-Gal4)*^*+/Tg*^*; Tg(UAS:EGFP)*^*+/Tg*^ larvae from the same clutch were used. At 5 dpf, larvae were collected and fixed in 4% paraformaldehyde (PFA) in PBS at 4°C for overnight. Larvae were then washed with PBS for five times and then embedded in 1.2% agarose and 5% sucrose in PBS. Agarose blocks were trimmed by a razor and then incubated in PBS containing 30% sucrose at 4°C for overnight. After replacement with 30% sucrose solution, blocks were frozen on dry ice and stored at -80°C until cryosectioning. Larvae were transversely sectioned (thickness = 16 μm) using a Leica CM 3050 S (Leica) and sections posterior to the swimming bladder were collected (one section per individual). Cryosections were adhered on a MAS-GP typeA-coated slide glass (MATSUNAMI GLASS Ind., LTD) and air-dried at room temperature for 30 min. Sections were rehydrated by PBS at room temperature for 30 min, and then permeabilized and blocked with 5% normal goat serum in PBS supplemented with 0.5% TritonX-100 (0.5% PBT) for 1 h. Sections were then incubated with the following primary antibodies diluted in 5% normal goat serum in 0.5% PBT at 4°C for overnight: rabbit anti-phosphorylated-Histone H3 (Ser10) (pH3) (EMD Millipore, 06-570; 1:100 dilution) and rabbit anti-E-cadherin (cdh1) (Gene Tex, GTX125890; 1:100 dilution). Sections were washed with 0.5% PBT and then incubated with secondary antibody, Alexa Fluor 568 conjugating anti-Rabbit IgG (Life Technology; 1:400 dilution), at room temperature for 1 h. Sections were washed with 0.5% PBT and then mounted with ProLong Gold Antifade Mount with DAPI (Thermo Fisher SCIENTIFIC). Fluorescent images were taken by Nikon A1R confocal laser microscope (Nikon).

### BrdU incorporation, cryosectioning and fluorescent immunohistochemistry

*Tg(5×UAS:EGFP-P2A-kras*^*G12D*^)^*+/Tg*^*; Tg(pInt-Gal4)*^*+/Tg*^ and *Tg*(*pInt-Gal4)*^*+/Tg*^*;Tg(UAS:EGFP)*^*+/Tg*^ larvae from the same clutch were used. BrdU incorporation experiments were performed essentially as described previously (Takada et al., 2010). At 4 dpf, 20 larvae were transferred into egg water containing 0.5 mM Bromodeoxyuridine (BrdU; nakalai tesque) and incubated for 24 h. At 5 dpf, larvae were rinsed with egg water and then fixed with 4% PFA in PBS. Agarose embedding and cryosectioning were performed as described above. After rehydration of cryosections by PBS, sections were treated with 2N hydrochloric acid to denature DNA at room temperature for 1 h and then washed with PBS. Blocking and antibody treatment were performed as described above. Primary antibodies, mouse anti-BrdU antibody (The Developmental Studies Hybridoma Bank, G3G4; 1:500 dilution) and rabbit anti-GFP antibody (MBL, 598; 1:500 dilution), and secondary antibodies, Alexa Fluor 568 conjugating anti-Mouse IgG (Life technologies; 1:500 dilution) and Alexa Fluor 488 conjugating anti-Rabbit IgG (Life technologies; 1:500 dilution) were used. Sections were counterstained with Hoechst33342 (Life technologies; 1:2000 dilution) and mounted with 80% glycerol in PBS. Fluorescent images were taken by Nikon A1R confocal laser microscope (Nikon).

### Whole mount fluorescent immunohistochemistry

*Tg(5×UAS:EGFP-P2A-kras*^*G12D*^)^*+/Tg*^*; Tg(pInt-Gal4)*^*+/Tg*^ and *Tg*(*pInt-Gal4)*^*+/Tg*^*; Tg(UAS:EGFP)*^*+/Tg*^ larvae from the same clutch were used. At 5 dpf, larvae ware fixed in 4% PFA in PBS at 4°C for overnight. Larvae were washed with PBS for five times and treated with 3% hydrogen peroxide in 0.5% sodium hydride at room temperature to bleach pigments. After removing pigments, larvae ware washed with PBS and then transferred into methanol, and stored at -30°C until staining. Larvae were washed with 0.5% PBT for 5 times. Permeabilization was performed by treating samples with distilled water for 5 min and then with cold acetone (-30°C) for 5 min. Larvae were washed with 0.5% PBT for 3 times and blocked with 5% goat serum in 0.5% PBT for 1 h. Larvae were incubated with rabbit anti-Lysozyme (Lyz) antibody (AnaSpec, AS-55633; 1:200 dilution) diluted in 5% normal goat serum in 0.5% PBT at 4°C for overnight. After washing with 0.5% PBT, samples were incubated with secondary antibody, Alexa Fluor 568 conjugating anti-Rabbit IgG (Life Technologies; 1:200 dilution) at room temperature for 1h. Larvae were counterstained with Hoechst33342 (Life Technologies; 1:2000 dilution) and mounted with PBS containing 80% glycerol. Fluorescent images were taken by Nikon A1R confocal laser microscope (Nikon).

### Paraffin sectioning and HE staining

Zebrafish larvae were fixed in 4% PFA in PBS at 4°C for overnight. Fixed larvae were dehydrated by a series of diluted ethanol (70, 80, 90, 99.5 and 100%) and xylene. Paraffin filtration was performed at 65°C for overnight, and then samples were embedded in paraffin at room temperature. Paraffin sectioning (thickness = 5 μm) was performed with HM 340E Rotary Microtome (Thermo Fisher SCIENTIFIC). Sections posterior to the pancreas were collected and deparaffinized by xylene and ethanol treatments, and then stained with Mayer’s Hematoxylin and eosinY (Wako Pure Chemical Industries). Images were taken using Nikon ECLIPSE Ni-E (Nikon).

### Imaging of neutrophils using Tg(lyz:EGFP)

The sibling controls and tumor-bearing fish carrying *Tg(lyz:EGFP)* and *Tg(fabp10a:mCherry)* or *Tg(fabp10a:mCherry-P2A-cyp7a1)* were obtained from the same clutch. At 7 dpf, larvae were given an anesthetic by 0.008% MS-222 and mounted in 1% NuSieve GTG Agarose (Lonza) in egg water. Fluorescent images of the left side of the liver were obtained using Nikon A1R confocal laser microscope (Nikon). Liver size was measured using ImageJ software (Schneider et al., 2012). The number of neutrophils overlapping with mCherry signals (i.e the liver) were manually counted using ImageJ software in all sections containing the liver (6 μm interval).

### Body length measurement

*Tg(5×UAS:EGFP-P2A-kras*^*G12D*^)^*+/Tg*^*; Tg(pInt-Gal4)*^*+/Tg*^ and the sibling controls were obtained from the same clutch. Embryos and larvae were reared in a plastic petri dish in the presence of egg water without foods. At 7 or 9 dpf, zebrafish larvae were given an anesthetic by 0.008% MS-222 and phenotyped into tumor-bearing fish and the sibling controls. Larvae were placed on the bottom of a plastic petri dish and lateral view images were taken by Leica DFC310 FX. Lengths of the lateral side views were measured by ImageJ software.

### Oil Red O staining

Oil Red O was purchased from Wako Pure Chemical Industries (Wako) and the experiments were performed essentially as described previously (Kim et al., 2013), except that we did not perform a rinse with 2-propanol after Oil Red O treatment.

### Survival assay

Twenty larvae of the sibling controls and tumor-bearing fish were reared in a tank from 7 to 14 dpf. Larvae were fed the Hikari Lab 130 food (KYORIN). The numbers of live and dead fish were counted everyday.

### RNA-seq and Bioinformatic analysis

RNA-seq analyses were performed as described previously (Kawaoka et al., 2013; Suzuki et al., 2014). 7 dpf larvae were dissected under a microscope. The liver, intestine, and the rest part of the body from 20 ∼ 30 of tumor-bearing fish or the sibling controls were pooled and RNA-extracted. Pooling multiple fish for preparing sequencing libraries was important to obtain sufficient amount of high-quality RNAs and to minimize the risk to obtain outliers that cannot represent the clutch used. The obtained gene list with reads per million per a kilobase (RPKM) scores were shown in Table S2. To identify differentially expressed genes (DEGs), we first focused on the well-annotated protein-coding genes. RPKM scores were used to calculate the ratio tumor/control. In this calculation, 1 was added to all RPKM scores to ignore the scores below “1", and to make analyses more stringent. Recognizing that our dissection cannot prevent cross-contamination, genes showing more than 0.8-fold-enrichment and > 0 RPKM in the tissue of interest were further considered. The obtained ratios were used to sort genes to find potential DEGs. As an initial screening to identify reliable DEGs, we focused on a set of genes showing more than 3-fold changes in the RNA-seq experiments. Considering possible differences among clutches, the RNA-seq experiment was followed by qPCR validation with samples prepared from different clutches. Thus, the RNA-seq experiment functioned as a screening to identify DEGs. Data visualization was done mostly using ggplot2 (http://ggplot2.org/). In main figures, we show genes consistently validated by qPCR. In our experience with our dataset, the validation rate was high for genes with more than 3-fold changes in the intestine-derived samples. In the liver and rest part of the body, “3-fold criteria” was not enough to obtain a high validation rate (i.e. genes showing more than 3-fold changes such as *pklr* failed to be validated by qPCR (data not shown)). Used in-house R scripts are all available upon request. RNA-seq data published in the present study have been deposited under the accession number of DRA005199 in DDBJ (DNA Data Bank of Japan).

### Metabolite measurement

For measuring total bile acids, single zebrafish larva was homogenized in 500 μL of chloroform: methanol (1:1) solution to extract total lipids. Samples were centrifuged at 20,000 g for 20 min at RT. Supernatants were collected and evaporated. Dried samples were dissolved in 75 μL of R1 reagent of total Bile Acids Assay Kit (DIAZYME, CA, USA), and then 25 μL of R2 reagent were added. Absorbance at 405 nm was measured by Multiskan GO (Thermo Fisher SCIENTIFIC). Standard curve was generated using dilution series of standard bile acids. For cholesterol measurements, single zebrafish larva was homogenized in 500 μL of chloroform: methanol (2:1) solution to extract total lipids. Samples were centrifuged at 20,000 g for 20 min at RT. Supernatants were collected and evaporated. Dried samples were dissolved in 100 μL of the assay reagent of the WAKO cholesterol E-test (Wako Pure Chemical Industries, Osaka, Japan). Absorbance at 405 nm was measured by Multiskan GO. Standard curve was generated using dilution series of standard cholesterol. The obtained data were shown as box plots generated using ggplot2. Used in-house R scripts are all available upon request.

### Statistics and sample size determination

The values of the bar graphs are expressed as average ± s.e.m. The error bars (s.e.m.) shown for all results were derived from biological replicates. Significant differences between two groups were examined using one or two-tailed, unpaired *t*-test. One-tailed test was chosen when we had hypothesis regarding direction of changes (increased or decreased) in experiments. Statistical significance is assumed if *p* < 0.05. The sample size was not pre-determined and chosen as follows. First, the number of animals was minimized as much as possible in light of animal ethics. Second, against effect size estimated in each experiment, ≥ 80%-90% power was favored. Third, in most cases, n ≥ 5 was set as a threshold according to the previous reports (Krzywinski and Altman, 2014). For analyzing the growth defect phenotype, with the estimated size effect (around 0.98-0.99 fold), larger sample size (e.g. n >50) was preferred to obtain appropriate statistical power. We did not find apparently abnormal distribution throughout the study except the controls in Figs. 2S and S2J, where the majority of controls exhibit 0. No data exclusion was performed.

## Acknowledgements

We thank Dr. Thomas. N. Sato (T.N.S), the director of The TNS BioMEC-X Laboratories, ATR, and JST ERATO Sato Live Bio-forecasting project, for supporting all aspects of the study. We thank Tomoko Kuroda, Tomoko Ninomiya, Satsuki Endo, Fumihiko Sagawa, Hitomi Anabuki, Satoshi Kozawa, Terumi Horiuchi, and Kiyomi Imamura for technical assistance. We thank Ryoko Takahashi, Erika Kojima, and Toshiya Morie for administrative assistance. We thank Dr. Norio Takada and Dr. Sa Kan Yoo for providing valuable suggestions on zebrafish usage. We thank Dr. Yoichiro Tamori for helpful discussion on tumor classification. We are thankful to Dr. Pieter Bas Kwak and Dr. Bryce Nelson for critically reading the manuscript. Thomas. N. Sato and the members of the TNS BioMEC-X Laboratories provided the insightful comments on the manuscript. This work was supported by JST ERATO (T.N.S; JPMJER1303) and Uehara Memorial Foundation Research Grant (S.K). This work was supported by National BioResource Project from AMED (K.K.), JSPS KAKENHI Grant Number JP15H02370 (K.K.). The funders had no role in study design, data collection and analysis, decision to publish, or preparation of the manuscript.

## Supplementary Figure Legends

**Figure S1 *aInt-Gal4* is expressed in the epidermis at 2 dpf**

Bright filed (A, B) and EGFP (C, D) images of the sibling controls and *EGFP-kras*^*G12D*^-expressing fish driven by gSAIzGFFM103B (*aInt-Gal4*) at 2 dpf. White arrows indicate *EGFP-kras*^*G12D*^-expressing cells. Scale bar represents 500 μm.

**Figure S2 Characterization of the pInt-Gal4-driven tumor model**

(A) A gel image of genotyping of tumor-bearing fish. Band sizes detecting *Tg(5×UAS-EGFP-P2A-kras*^*G12D*^), *Tg*(*pInt-Gal4)* and *tp53* are 701 bp, 345 bp and 88 bp, respectively. *tp53* locus is used as a PCR control. M, DNA ladder marker: W, wild type fish: G, parental *Tg(pInt-Gal4)* line: K, parental *Tg(5×UAS-EGFP-P2A-kras*^*G12D*^) line.

(B)-(I) Representative images of fluorescent immunohistochemistry for BrdU and EGFP in intestine sections of the sibling controls and tumor bearing fish at 5 dpf. BrdU (B, C), Hoechst33342 (D, E) and EGFP (F, G) images are shown. In the merged images (H, I), BrdU, Hoechst33342 and EGFP signals are shown in red, blue and green, respectively. White arrows indicate intestinal cells positive for BrdU, Hoechst33342, and EGFP. Scale bar indicates 100 μm.

(J) The number of BrdU and EGFP positive intestinal cells. The number of BrdU and EGFP positive cells was counted from single section per individual fish. The data harbors 10 and 9 biological replicates from the sibling controls and tumor-bearing fish, respectively. Error bars represent ± s.e.m.

**Figure S3 The intestinal rumen is not completely disrupted in tumor-bearing fish** Representative images of the sibling controls (A) and tumor-bearing fish (B) at 9 dpf in the presence of foods in the intestine are shown. Scale bar represents 500 μm.

**Figure S4 The intestinal tumor causes systemic growth defects in multiple independent clutches**

Body length data from three individual clutches at 7 dpf (A) and 9 dpf (B) are presented. Error bars represent ± s.e.m. The numbers within the bars indicate the number of biological replicates in each clutch.

**Figure S5 The effects of the intestinal tumor on whole-organismal gene expression and cholesterol-BAs metabolism**

(A) Measurement for systemic cholesterol levels at 7 and 9 dpf (n = 12 for 7 dpf and 16-19 for 9 dpf). Statistical significance was tested using student’s *t*-test (unpaired, one-tailed).

(B)-(C) Gene expression levels of *ggt1* (B) and *mpx* (C) in the intestine at 9 dpf are shown. The scores are normalized by expression of *rpl13a*. The data harbors 5 biological replicates, each containing 5 fish. Error bars represent ± s.e.m. Statistical significance was tested using student’s *t*-test (unpaired, one-tailed).

**Figure S6 Correlation between body length and total BAs levels**

Correlation between body length and total bile acids levels at 9 dpf (n = 20 per a group).

**Table S1 The primers and DNA sequences used in the study**

**Table S2 Raw RPKM scores determined by RNA-seq analysis**

**Table S3 Calculation for sample enrichments**

**Table S4 The list of 8261 liver enriched genes (Liver to body > 0.8, Control-liver > 0)**

**Table S5 The list of 7294 body-enriched genes (Body to liver > 0.8, Body to intestine > 0.8m Control-body > 0)**

**Table S6 The list of 8002 intestine-enriched genes (Intestine to body > 0.8, Control-intestine > 0)**

## References

Asakawa, K. and Kawakami, K. (2008). Targeted gene expression by the Gal4— UAS system in zebrafish. Dev Growth Differ 50, 391–9.

Asakawa, K., Suster, M. L., Mizusawa, K., Nagayoshi, S., Kotani, T., Urasaki, A., Kishimoto, Y., Hibi, M. and Kawakami, K. (2008). Genetic dissection of neural circuits by Tol2 transposon—mediated Gal4 gene and enhancer trapping in zebrafish. Proc Natl Acad Sci U S A 105, 1255–60.

Baxter, R. C. (2014). IGF binding proteins in cancer: mechanistic and clinical insights. Nat Rev Cancer 14, 329–41.

Bonetto, A., Rupert, J. E., Barreto, R. and Zimmers, T. A. (2016). The Colon— 26 Carcinoma Tumor—bearing Mouse as a Model for the Study of Cancer Cachexia. J Vis Exp.

Burnicka—Turek, O., Mohamed, B. A., Shirneshan, K., Thanasupawat, T., Hombach—Klonisch, S., Klonisch, T. and Adham, I. M. (2012). INSL5—deficient mice display an alteration in glucose homeostasis and an impaired fertility. Endocrinology 153, 4655–65.

Colombani, J., Andersen, D. S., Boulan, L., Boone, E., Romero, N., Virolle, V., Texada, M. and Leopold, P. (2015). Drosophila Lgr3 Couples Organ Growth with Maturation and Ensures Developmental Stability. Curr Biol 25, 2723–9.

Colombani, J., Andersen, D. S. and Leopold, P. (2012). Secreted peptide Dilp8 coordinates Drosophila tissue growth with developmental timing. Science 336, 582–5.

Das, S. K., Eder, S., Schauer, S., Diwoky, C., Temmel, H., Guertl, B., Gorkiewicz, G., Tamilarasan, K. P., Kumari, P., Trauner, M. et al. (2011). Adipose triglyceride lipase contributes to cancer—associated cachexia. Science 333, 233–8.

Degirolamo, C., Sabba, C. and Moschetta, A. (2016). Therapeutic potential of the endocrine fibroblast growth factors FGF19, FGF21 and FGF23. Nat Rev Drug Discov 15, 51–69.

Egeblad, M., Nakasone, E. S. and Werb, Z. (2010). Tumors as organs: complex tissues that interface with the entire organism. Dev Cell 18, 884–901.

Evason, K. J., Francisco, M. T., Juric, V., Balakrishnan, S., Lopez Pazmino Mdel, P., Gordan, J. D., Kakar, S., Spitsbergen, J., Goga, A. and Stainier, D. Y. (2015). Identification of Chemical Inhibitors of beta—Catenin—Driven Liver Tumorigenesis in Zebrafish. PLoS Genet 11, e1005305.

Fearon, K. C., Glass, D. J. and Guttridge, D. C. (2012). Cancer cachexia: mediators, signaling, and metabolic pathways. Cell Metab 16, 153–66.

Feng, Y., Renshaw, S. and Martin, P. (2012). Live imaging of tumor initiation in zebrafish larvae reveals a trophic role for leukocyte—derived PGE(2). Curr Biol 22, 1253–9.

Feng, Y., Santoriello, C., Mione, M., Hurlstone, A. and Martin, P. (2010). Live imaging of innate immune cell sensing of transformed cells in zebrafish larvae: parallels between tumor initiation and wound inflammation. PLoS Biol 8, e1000562.

Figueroa—Clarevega, A. and Bilder, D. (2015). Malignant Drosophila tumors interrupt insulin signaling to induce cachexia—like wasting. Dev Cell 33, 47–55.

Garelli, A., Gontijo, A. M., Miguela, V., Caparros, E. and Dominguez, M. (2012). Imaginal discs secrete insulin—like peptide 8 to mediate plasticity of growth and maturation. Science 336, 579–82.

Garelli, A., Heredia, F., Casimiro, A. P., Macedo, A., Nunes, C., Garcez, M., Dias, A. R., Volonte, Y. A., Uhlmann, T., Caparros, E. et al. (2015). Dilp8 requires the neuronal relaxin receptor Lgr3 to couple growth to developmental timing. Nat Commun 6, 8732.

Grosse, J., Heffron, H., Burling, K., Akhter Hossain, M., Habib, A. M., Rogers, G. J., Richards, P., Larder, R., Rimmington, D., Adriaenssens, A. A. et al. (2014). Insulin— like peptide 5 is an orexigenic gastrointestinal hormone. Proc Natl Acad Sci U S A 111, 11133–8.

Hanahan, D. and Weinberg, R. A. (2011). Hallmarks of cancer: the next generation. Cell 144, 646–74.

Her, G. M., Chiang, C. C., Chen, W. Y. and Wu, J. L. (2003). In vivo studies of liver—type fatty acid binding protein (L—FABP) gene expression in liver of transgenic zebrafish (Danio rerio). FEBS Lett 538, 125–33.

Hojo, H., Enya, S., Arai, M., Suzuki, Y., Nojiri, T., Kangawa, K., Koyama, S. and Kawaoka, S. (2017). Remote reprogramming of hepatic circadian transcriptome by breast cancer. Oncotarget.

Katsuyama, T., Comoglio, F., Seimiya, M., Cabuy, E. and Paro, R. (2015). During Drosophila disc regeneration, JAK/STAT coordinates cell proliferation with Dilp8— mediated developmental delay. Proc Natl Acad Sci U S A 112, E2327—36.

Kaufman, C. K., Mosimann, C., Fan, Z. P., Yang, S., Thomas, A. J., Ablain, J., Tan, J. L., Fogley, R. D., van Rooijen, E., Hagedorn, E. J. et al. (2016). A zebrafish melanoma model reveals emergence of neural crest identity during melanoma initiation. Science 351, aad2197.

Kawakami, K. (2004). Transgenesis and gene trap methods in zebrafish by using the Tol2 transposable element. Methods Cell Biol 77, 201–22.

Kawakami, K., Asakawa, K., Hibi, M., Itoh, M., Muto, A. and Wada, H. (2016). Gal4 Driver Transgenic Zebrafish: Powerful Tools to Study Developmental Biology, Organogenesis, and Neuroscience. Adv Genet 95, 65–87.

Kawakami, K., Koga, A., Hori, H. and Shima, A. (1998). Excision of the tol2 transposable element of the medaka fish, Oryzias latipes, in zebrafish, Danio rerio. Gene 225, 17–22.

Kawaoka, S., Hara, K., Shoji, K., Kobayashi, M., Shimada, T., Sugano, S., Tomari, Y., Suzuki, Y. and Katsuma, S. (2013). The comprehensive epigenome map of piRNA clusters. Nucleic Acids Res 41, 1581–90.

Kim, J. H., Lee, S. R., Li, L. H., Park, H. J., Park, J. H., Lee, K. Y., Kim, M. K., Shin, B. A. and Choi, S. Y. (2011). High cleavage efficiency of a 2A peptide derived from porcine teschovirus—1 in human cell lines, zebrafish and mice. PLoS One 6, e18556.

Kim, S. H., Scott, S. A., Bennett, M. J., Carson, R. P., Fessel, J., Brown, H. A. and Ess, K. C. (2013). Multi—organ abnormalities and mTORC1 activation in zebrafish model of multiple acyl—CoA dehydrogenase deficiency. PLoS Genet 9, e1003563.

Kir, S., Komaba, H., Garcia, A. P., Economopoulos, K. P., Liu, W., Lanske, B., Hodin, R. A. and Spiegelman, B. M. (2016). PTH/PTHrP Receptor Mediates Cachexia in Models of Kidney Failure and Cancer. Cell Metab 23, 315–23.

Kir, S., White, J. P., Kleiner, S., Kazak, L., Cohen, P., Baracos, V. E. and Spiegelman, B. M. (2014). Tumour—derived PTH—related protein triggers adipose tissue browning and cancer cachexia. Nature 513, 100–4.

Kitaguchi, T., Kawakami, K. and Kawahara, A. (2009). Transcriptional regulation of a myeloid—lineage specific gene lysozyme C during zebrafish myelopoiesis. Mech Dev 126, 314–23.

Krzywinski, M. and Altman, N. (2014). Points of significance: Comparing samples—part I. Nat Methods 11, 215–6.

Kuipers, F., Bloks, V. W. and Groen, A. K. (2014). Beyond intestinal soap——bile acids in metabolic control. Nat Rev Endocrinol 10, 488–98.

Kwon, Y., Song, W., Droujinine, I. A., Hu, Y., Asara, J. M. and Perrimon, N. (2015). Systemic organ wasting induced by localized expression of the secreted insulin/IGF antagonist ImpL2. Dev Cell 33, 36–46.

Lieffers, J. R., Mourtzakis, M., Hall, K. D., McCargar, L. J., Prado, C. M. and Baracos, V. E. (2009). A viscerally driven cachexia syndrome in patients with advanced colorectal cancer: contributions of organ and tumor mass to whole—body energy demands. Am J Clin Nutr 89, 1173–9.

Lister, J. A., Capper, A., Zeng, Z., Mathers, M. E., Richardson, J., Paranthaman, K., Jackson, I. J. and Patton, E. E. (2014). A conditional zebrafish MITF mutation reveals MITF levels are critical for melanoma promotion vs. regression in vivo. J Invest Dermatol 134, 133–40.

Liu, H., Pathak, P., Boehme, S. and Chiang, J. Y. (2016). Cholesterol 7alpha— hydroxylase protects the liver from inflammation and fibrosis by maintaining cholesterol homeostasis. J Lipid Res 57, 1831–1844.

Lobert, V. H., Mouradov, D. and Heath, J. K. (2016). Focusing the Spotlight on the Zebrafish Intestine to Illuminate Mechanisms of Colorectal Cancer. Adv Exp Med Biol 916, 411–37.

Mayrhofer, M., Gourain, V., Reischl, M., Affaticati, P., Jenett, A., Joly, J. S., Benelli, M., Demichelis, F., Poliani, P. L., Sieger, D. et al. (2017). A novel brain tumour model in zebrafish reveals the role of YAP activation in MAPK— and PI3K—induced malignant growth. Dis Model Mech 10, 15–28.

McAllister, S. S. and Weinberg, R. A. (2014). The tumour—induced systemic environment as a critical regulator of cancer progression and metastasis. Nat Cell Biol 16, 717–27.

Mione, M. and Zon, L. I. (2012). Cancer and inflammation: an aspirin a day keeps the cancer at bay. Curr Biol 22, R522–5.

Narsale, A. A., Enos, R. T., Puppa, M. J., Chatterjee, S., Murphy, E. A., Fayad, R., Pena, M. O., Durstine, J. L. and Carson, J. A. (2015). Liver inflammation and metabolic signaling in ApcMin/+ mice: the role of cachexia progression. PLoS One 10, e0119888.

Okin, D. and Medzhitov, R. (2016). The Effect of Sustained Inflammation on Hepatic Mevalonate Pathway Results in Hyperglycemia. Cell 165, 343–56.

Omae, M., Takada, N., Yamamoto, S., Nakajima, H. and Sato, T. N. (2013). Identification of inter—organ vascular network: vessels bridging between organs. PLoS One 8, e65720.

Owusu—Ansah, E. and Perrimon, N. (2015). Stress signaling between organs in metazoa. Annu Rev Cell Dev Biol 31, 497–522.

Patton, E. E. (2012). Live imaging in zebrafish reveals neu(trophil) insight into the metastatic niche. J Pathol 227, 381–4.

Robison, L. L. and Hudson, M. M. (2014). Survivors of childhood and adolescent cancer: life—long risks and responsibilities. Nat Rev Cancer 14, 61–70.

Rose, S. R., Horne, V. E., Howell, J., Lawson, S. A., Rutter, M. M., Trotman, G. E. and Corathers, S. D. (2016). Late endocrine effects of childhood cancer. Nat Rev Endocrinol 12, 319–36.

Santoriello, C., Gennaro, E., Anelli, V., Distel, M., Kelly, A., Koster, R. W., Hurlstone, A. and Mione, M. (2010). Kita driven expression of oncogenic HRAS leads to early onset and highly penetrant melanoma in zebrafish. PLoS One 5, e15170.

Schneider, C. A., Rasband, W. S. and Eliceiri, K. W. (2012). NIH Image to ImageJ: 25 years of image analysis. Nat Methods 9, 671–5.

Schubbert, S., Shannon, K. and Bollag, G. (2007). Hyperactive Ras in developmental disorders and cancer. Nat Rev Cancer 7, 295–308.

Suzuki, A., Makinoshima, H., Wakaguri, H., Esumi, H., Sugano, S., Kohno, T., Tsuchihara, K. and Suzuki, Y. (2014). Aberrant transcriptional regulations in cancers: genome, transcriptome and epigenome analysis of lung adenocarcinoma cell lines. Nucleic Acids Res 42, 13557–72.

Takada, N., Kucenas, S. and Appel, B. (2010). Sox10 is necessary for oligodendrocyte survival following axon wrapping. Glia 58, 996–1006.

Thermes, V., Grabher, C., Ristoratore, F., Bourrat, F., Choulika, A., Wittbrodt, J. and Joly, J. S. (2002). I—SceI meganuclease mediates highly efficient transgenesis in fish. Mech Dev 118, 91–8.

Thomas, C., Pellicciari, R., Pruzanski, M., Auwerx, J. and Schoonjans, K. (2008). Targeting bile—acid signalling for metabolic diseases. Nat Rev Drug Discov 7, 678–93.

Vallejo, D. M., Juarez—Carreno, S., Bolivar, J., Morante, J. and Dominguez, M. (2015). A brain circuit that synchronizes growth and maturation revealed through Dilp8 binding to Lgr3. Science 350, aac6767.

Wallace, K. N., Akhter, S., Smith, E. M., Lorent, K. and Pack, M. (2005). Intestinal growth and differentiation in zebrafish. Mech Dev 122, 157–73.

Wang, Z., Du, J., Lam, S. H., Mathavan, S., Matsudaira, P. and Gong, Z. (2010). Morphological and molecular evidence for functional organization along the rostrocaudal axis of the adult zebrafish intestine. BMC Genomics 11, 392.

White, R., Rose, K. and Zon, L. (2013). Zebrafish cancer: the state of the art and the path forward. Nat Rev Cancer 13, 624–36.

White, R. M., Cech, J., Ratanasirintrawoot, S., Lin, C. Y., Rahl, P. B., Burke, C. J., Langdon, E., Tomlinson, M. L., Mosher, J., Kaufman, C. et al. (2011). DHODH modulates transcriptional elongation in the neural crest and melanoma. Nature 471, 518–22.

White, R. M., Sessa, A., Burke, C., Bowman, T., LeBlanc, J., Ceol, C., Bourque, C., Dovey, M., Goessling, W., Burns, C. E. et al. (2008). Transparent adult zebrafish as a tool for in vivo transplantation analysis. Cell Stem Cell 2, 183–9.

